# Myosin inhibitor reverses hypertrophic cardiomyopathy in pediatric iPSC-cardiomyocytes to mirror variant correction

**DOI:** 10.1101/2023.04.14.536782

**Authors:** Caroline Kinnear, Abdelrahman Said, Guoliang Meng, Yimu Zhao, Erika Y Wang, Neha Parmar, Wei Wei, Filio Billia, Craig A Simmons, Milica Radisic, James Ellis, Seema Mital

## Abstract

Hypertrophic cardiomyopathy (HCM) is mainly caused by sarcomere gene variants in MYH7 and MYBPC3. Targeted drugs like myosin ATPase inhibitors have shown efficacy in adult HCM but have not been evaluated in children. We generated iPSC-cardiomyocytes (CMs) from four children with HCM harboring variants in *MYH7* (*V606M; R453C)* or *MYBPC3 (G148R; P955fs and TNNI3_A157P*), variant-corrected controls, and a healthy individual. All CMs showed hypertrophy and sarcomere disorganization. All 3 single variant CMs showed higher contractility, slower relaxation, higher calcium transients and higher ATPase activity. Only *MYH7* variant CMs showed stronger myosin-actinin binding. Targeted myosin ATPase inhibitor showed complete rescue of the phenotype in affected CMs and in cardiac Biowires to mirror isogenic controls. The response was stronger compared to verapamil or metoprolol, highlighting the need for clinical trials of myosin targeted therapy in pediatric HCM patients. The phenotype and response to drug therapy are influenced by the underlying genotype.

## INTRODUCTION

Hypertrophic cardiomyopathy (HCM), characterized by abnormal ventricular thickening, is the most common form of cardiomyopathy and the leading cause of heart failure and sudden cardiac death in children.^1^ The hallmark of HCM is left ventricular hypertrophy, myocyte disarray and hyperdynamic contractility, with a subset of patients progressing to systolic and/or diastolic heart failure.^2; 3^

About 70% of HCM patients harbor heterozygous variants in one of two sarcomere genes: *MYH7* that encodes myosin heavy chain β, and *MYBPC3* that encodes myosin-binding protein C.^4^ Both MHC-β and cMyBP-C are located in the thick filament of the sarcomere where myosin interacts with actin in a process coupled to the hydrolysis of ATP and governed by calcium fluxes, which drives myocardial force generation and contraction.^5^ Missense variants in *MYH7* and *MYBPC3* cause HCM due to incorporation of mutant myosins into the sarcomere. Besides missense variants, protein-truncating variants in *MYBPC3* result in a premature termination codon that drives nonsense-mediated decay of mRNA and haploinsufficiency of cMyBP-C to cause HCM.^6^

Current medical therapies include L-type calcium channel blockers and β-blockers.^7^ However, these treatments only offer symptomatic relief. They do not target the underlying genetic cause or mechanism of disease and therefore do not alter disease progression. A myosin ATPase inhibitor, mavacamten, was recently approved as a targeted drug for adults with HCM based on clinical trials that showed an improvement in ventricular hypertrophy, exercise capacity, biomarkers, and overall health status.^8^ Despite the majority of pediatric HCM cases being caused by *MYH7* and *MYBPC3* variants, a myosin inhibitor has not been studied in the pediatric population. Further, it is unclear if genotype influences drug responses in patients with HCM.

Human induced pluripotent stem cell (iPSC)-derived cardiomyocytes (CMs) have been used to model HCM and study drug responses, either through reprogramming of patient cells to iPSCs or through CRISPR-editing of variants into healthy control iPSCs.^9–21^ However, most studies were in adult patients, and most did not include isogenic controls for comparison. Previous studies have reported that up to 16% of pediatric patients with HCM harbor multiple variants,^22; 23^ but drug responses in this context have not been studied. We generated iPSC-derived CMs from pediatric patients with HCM caused by variants in *MYH7* or *MYBPC3*, confirmed that the variants were disease-causing by showing phenotypic rescue with variant-correction, and compared the relative efficacy of standard drugs versus a myosin-targeted drug in rescuing the abnormal phenotype across different variant types. We also generated cardiac Biowires in a subset of patients using these patient CMs to validate the HCM phenotype and drug effects in an engineered tissue model.

## RESULTS

### Patient clinical characteristics

Skin fibroblasts or lymphoblasts were reprogrammed from 4 patients with HCM recruited through our Heart Centre Biobank.^24^ Two patients harbored single missense variants in *MYH7* i.e., V606M (iPSC line 80) and R453C (iPSC line 81), one patient harbored a single missense variant in *MYBPC3* i.e., G148R (iPSC line 82), and another harbored two variants - a frameshift variant (P955fs) and an additional *TNNI3* missense variant (A157P) (iPSC line 83) (**Figure 1**). These variants were identified on clinical testing as well as on research whole genome sequencing and met the American College of Medical Genetics criteria for pathogenicity.^25^ Clinical characteristics of patients and family members are shown in **Table 1** and family pedigrees are shown in **Figure S1**. Patients 80, 81 and 83 had severe obstructive HCM requiring surgical septal reduction therapy, while patient 82 had mild concentric HCM that did not require surgical intervention.

**Figure 1.**
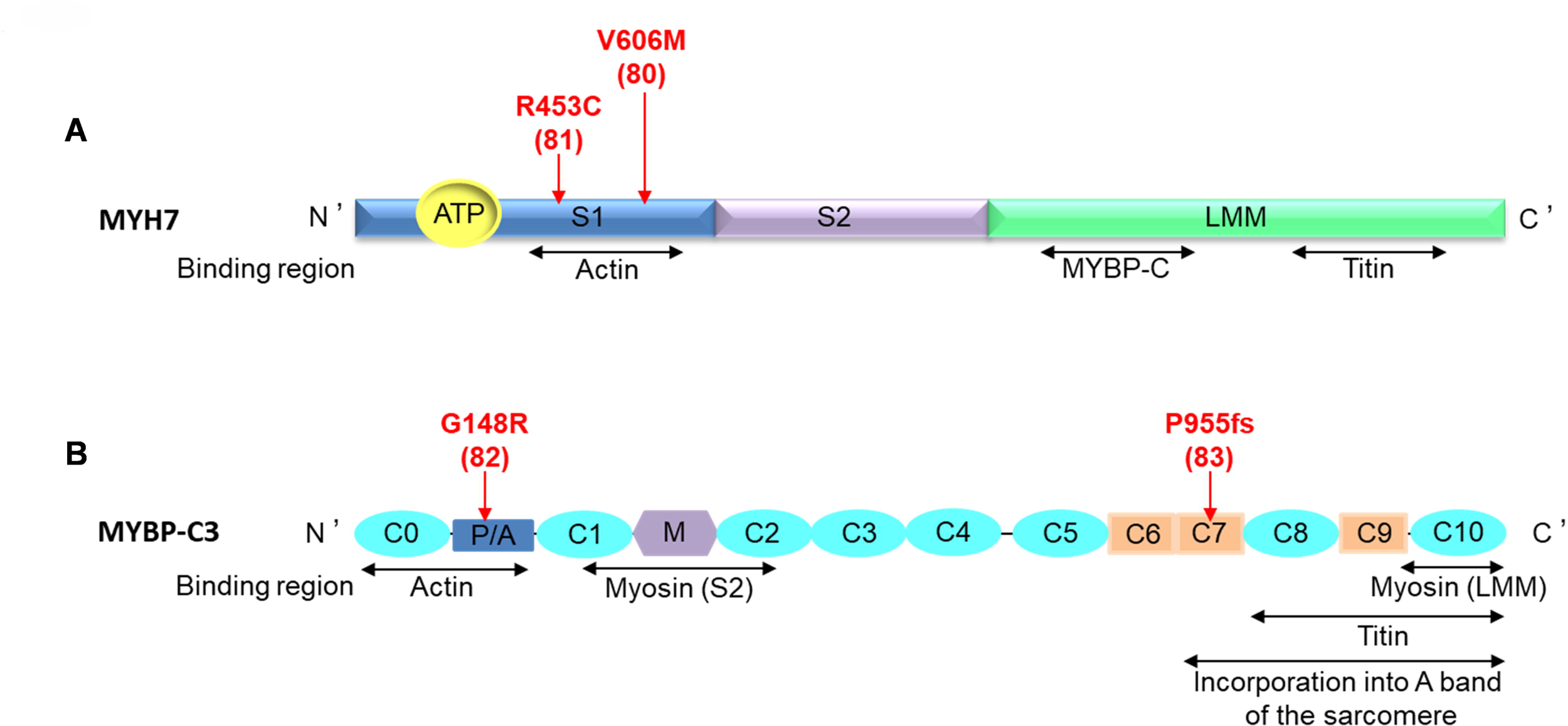
Location of variants in functional protein domains. **A.** Cardiac beta-myosin heavy chain encoded by the *MYH7* gene, consists of three regions: subfragment 1 (S1) or the head, subfragment 2 (S2) or the neck, and light meromyosin (LMM), also known as the tail. Arrows demonstrate the missense variants - V606M (patient 80) and R453C (patient 81) located in the region responsible for binding to actin. **B.** Cardiac myosin binding protein C3 (MYBP-C3). Ovals represent immunoglobulin-like domains and rectangles represent fibronectin type III domains. P/A is the proline/alanine region and M the phosphorylatable domain. The G148R missense variant (patient 82) is located in the actin binding region, while the P955fs protein-truncating variant (patient 83) is located in the region that incorporates into the A band of the sarcomere.

**Table 1.** Clinical characteristics of HCM patients.

| Patient ID (line ID) | Variants | Variant type | Age at diagnosis (years) | Sex | Family history of HCM | Age at reprogramm ing sample acquisition (years) | ECG at last follow-up | Echocardi ogram at last follow-up | Age at surgical myectomy (years) | Myocardial pathology |
| --- | --- | --- | --- | --- | --- | --- | --- | --- | --- | --- |
| 80 (80V) | *MYH7* V606M | Missense | 11.5 | M | Yes | 14.3 | LV hypertrophy ; QTc 484ms | Severe obstructive HCM | 14.3 | Myocyte hypertrophy and patchy interstitial fibrosis |
| 81 (81J) | *MYH7* R453C | Missense | 4.8 | M | No | 11 | LV hypertrophy with strain; QTc 409ms | Severe obstructive HCM | 6.4 & 11 | Myocyte hypertrophy and endocardial fibroelastosis |
| 82 (82U) | *MYBPC3* G148R | Missense | 0.9 | M | Yes | 2.1 | No LV hypertrophy ; QTc 433ms | Mild concentric LV hypertrophy | N/A | N/A |
| 83 (83M) | *MYBPC3* P955fs + *cTNNI3* A157V | *MYBPC3*: Frameshift<br>*cTNNI3*: Missense | 3.1 | M | No | 5.1 | LV hypertrophy ; QTc 457ms | Severe obstructive HCM | 5.1 | Myocyte hypertrophy, patchy myocardial fibrosis, myofiber disarray |

ECG, electrocardiogram; LV: Left ventricular; QTc, Corrected QT Interval; HCM, hypertrophic cardiomyopathy; N/A, not applicable

### Patient and variant-corrected iPSCs and CMs from HCM patients

We generated *MYH7* and *MYBPC3* variant lines after reprogramming using a Sendai virus vector carrying human transcription factors Oct4, Sox2, Kif4, and c-Myc.^26^ We confirmed normal karyotype and pluripotency of all iPSC lines (80, 81, 82, 83) (**Figures S2-S5**). CRISPR technology was used to generate myosin variant-corrected iPSCs as isogenic controls (80C, 81C, 82C, 83C). *TNNI3* variant was not corrected in patient 83. Variant correction and freedom from off-target events was confirmed (**Figure S2-S5**). A previously characterized iPSC line from a healthy individual free of cardiac variants (PGPC17) was used as a healthy control.^27^ All iPSC lines were cultured, differentiated into CMs, dissociated to single CMs at day 16, reseeded and maintained in iCell medium for phenotypic and functional assays. CM size and sarcomere organization were measured using immunofluorescence. Beat rate, contractility and relaxation were measured from days 30 to 40 using xCELLigence, a real-time cell analysis system. To delineate the mechanism of abnormal contractility in HCM, calcium (Ca2+) flux, ATPase activity and myosin-actinin co-immunoprecipitation assays were performed on days 38-42 when cells exhibited a more mature phenotype involving regular contractility and stable electrical signals. Drug treatments were performed on days 38-42. All experiments were done using 3 biological replicates i.e., 3 independent CM differentiations.

### Baseline phenotype of patient and variant-corrected CMs

#### Patient CMs showed hypertrophy and sarcomere disorganization that was rescued by variant-correction

We characterized the morphology of the CMs by co-immunostaining using antibodies against ventricular myosin light chain-2 (MLC2v) and sarcomeric α-actinin. Myocyte area was 1.4 to 2-fold larger in patient CMs (p<0.05 vs. PGPC17) and normalized after variant correction (p<0.05 vs. patient CMs) (**Figure 2A-B, Table S1**). Patient CMs showed reduced myosin-actinin alignment on line-scan analysis compared to PGPC17 (29%±3% vs 85%±5% respectively; p<0.05). Variant correction was associated with improved sarcomere organization with 72%±10% corrected CMs showing well-aligned myosin-actinin filaments (p<0.05 vs. patient CMs) (**Figure 2C-E, Table S1**). All patient CMs therefore recapitulated the morphological hallmark of HCM i.e., myocyte hypertrophy and sarcomere disorganization, and showed morphological rescue with variant-correction.

**Figure 2.**
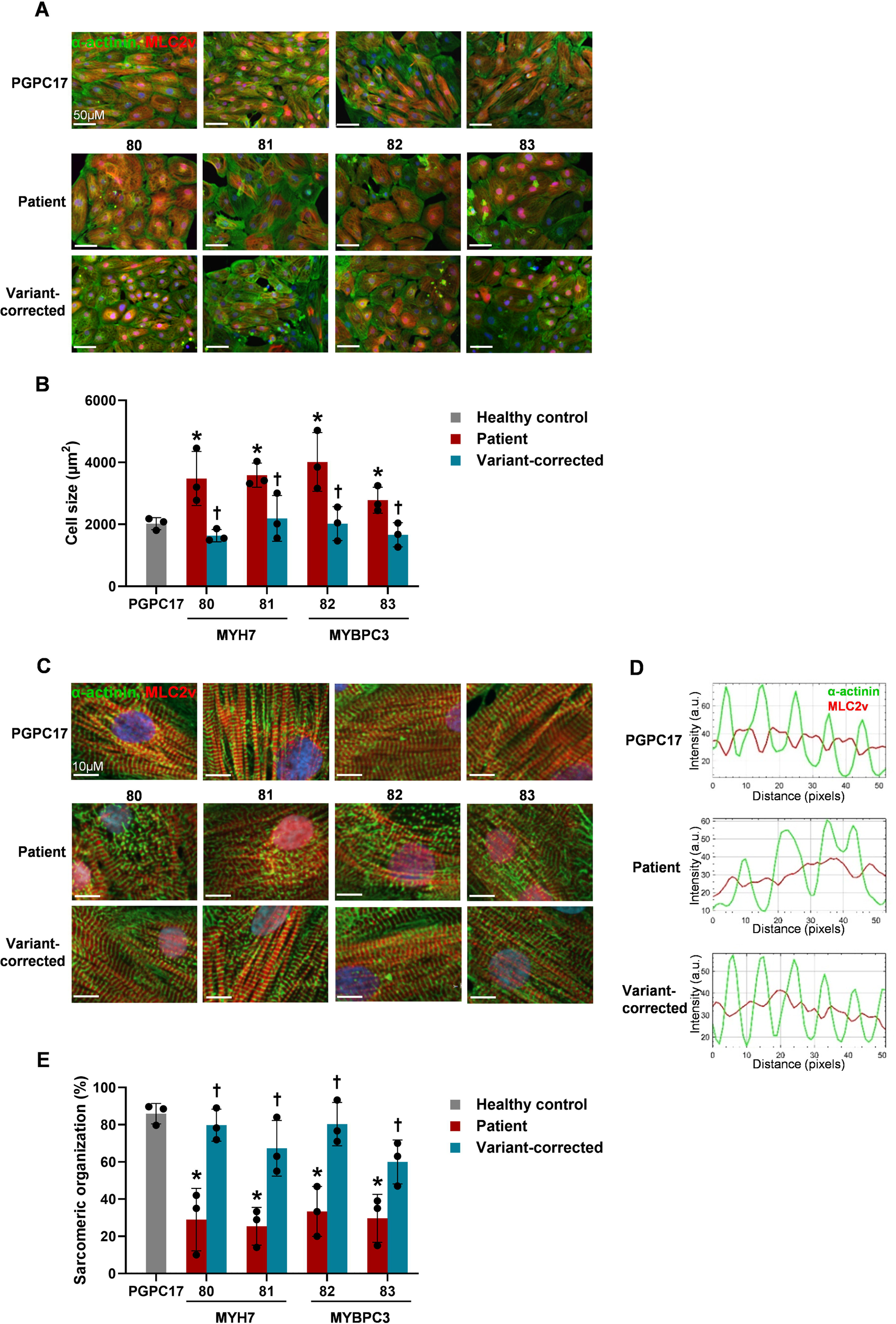
Cardiomyocyte size and sarcomere organization in healthy control, HCM patient and variant-corrected iPSC-CMs. Representative images and bar graphs showing results of immunofluorescence staining with α-actinin (green) and MLC2v (red) of healthy control, patient, and variant-corrected CMs. **A-B. Myocyte size:** All 4 patient lines showed increased myocyte size (3465±512µM^2^) compared to PGPC17 (2022±194µM^2^). Variant correction restored myocyte size to that seen in PCPC17 control line (1878±274µM^2^). **C-E. Sarcomere organization** was measured using line-scan analysis of α-actinin and MLC2v fluorescence intensity longitudinally throughout myofibers. PGPC17 control showed 85±5% CMs with well-aligned myosin actin filaments. The 4 patient lines (80, 81, 82, 83) showed a low percentage of CMs with organized sarcomeres (29±3%). CMs with sarcomere organization significantly improved in variant-corrected CMs to 72±10%. *p<0.05 patient vs. PGPC17, ^†^p<0.05 patient vs. variant-corrected. n=3 independent experiments, using 4 technical replicates for each experiment. Error bars represent standard deviation. 80, *MYH7* V606M; 81, *MYH7* R453C; 82, *MYBPC3* G148R; 83, *MYBPC3* P955fs and *TNNI3* A157V. CM, cardiomyocytes; MLC2v, ventricular myosin light chain 2.

#### Patient CMs showed abnormal contraction, relaxation and abnormal calcium transients

##### Contractility and relaxation

We assessed beat amplitude using xCELLigence impedance electrodes as a marker of myocyte contractility over the course of 10 days of differentiation from days 30-40 (**Figure 3A**). Spontaneous beat amplitude was higher in 80, 81 and 82 CMs (p<0.05 vs. PGPC17 and isogenic controls). Falling time, i.e., the time to achieve lowest impedance during the downstroke of each beat, was used as a marker of relaxation velocity. Falling time was longer in 80, 81 and 82 CMs indicating slower relaxation (p<0.05 vs. PGPC17 and isogenic controls). 83 CMs harboring both *MYBPC3* and *TNNI3* variants did not show contractile or relaxation abnormalities (**Figure 3B, C**). Spontaneous beat rate was similar in all patient CMs (**Figure 3D**). Absolute values are shown in **Table S1**.

**Figure 3.**
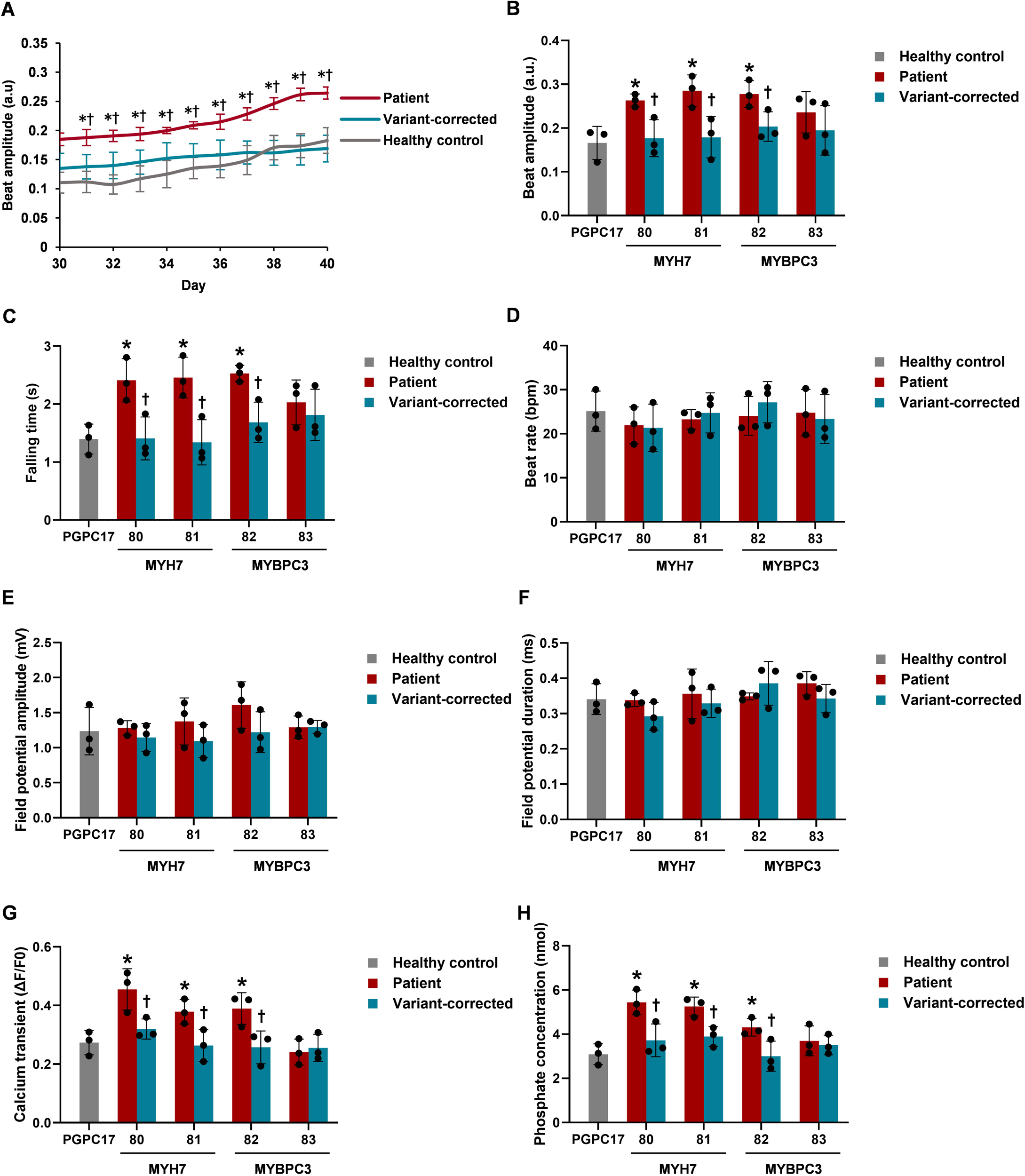
Contractile and electrophysiological phenotype of healthy control, HCM patient and variant-corrected iPSC-CMs. **A. Beat amplitude (serial recordings)** measured using the RTCA CardioECR xCELLigence system from day 30 to 40 was higher in patient 80 compared to PGPC17 healthy control and variant-corrected CMs. **B-H.** Box and whisker plots showing averaged values for functional parameters at day 40. **B. Beat amplitude** and **C. Falling time** were higher in 80, 81 and 82 patient CMs compared to PGPC17 control. Both abnormalities were rescued in the variant-corrected lines. The 83 CMs did not show higher beat amplitude or higher falling time compared to PGPC17. **D. Beat rate, E. Field potential amplitude**, and **F. Field potential duration** were not different between HCM patient CMs compared to PGPC17 control and variant-corrected CMs. **G. Calcium transients:** ΔF/F0, ratio of peak fluorescence intensity to baseline intensity, was higher in 80, 81, 82 compared to PGPC17 and variant-corrected CMs. **H. ATPase activity:** Phosphate concentration, a measure of ATPase activity, was higher in 80, 81 and 82 compared to PGPC17 and variant-corrected CMs. *p<0.05 patient vs. PGPC17, ^†^p<0.05 patient vs. corrected. n=3 independent experiments, using 4 technical replicates for each experiment. Error bars represent standard deviation. 80, *MYH7* V606M; 81, *MYH7* R453C; 82, *MYBPC3* G148R; 83, *MYBPC3* P955fs and *TNNI3* A157V. CM, cardiomyocytes; a.u., arbitrary unit; bpm, beat per minute.

##### Field potential amplitude and duration

The xCELLigence extracellular recordings were used to measure field potential amplitude (FPA) which is driven by the influx of sodium at the start of the cardiac action potential. The recordings also measured field potential duration (FPD), the time period between start of depolarization and peak or end of repolarization. FPA and FPD were not different between patient CMs compared to PGPC17 and variant-corrected controls (**Figure 3E-F, Table S1**).

##### Calcium transients

Ca2+ currents regulate cardiac excitation-contraction and were measured as intracellular Ca2+ transient amplitudes using Fluo-4 imaging. 80, 81 and 82 patient CMs showed higher Ca2+ transients (p<0.05 vs. PGPC17), that normalized in variant-corrected controls (p<0.05 vs. patient CMs). 83 CMs (*MYBPC3* P955fs/*TNNI3* A157P) did not show higher calcium transients (**Figure 3G, Table S1**).

Therefore, all single-variant *MYH7* and *MYBPC3* CMs showed hypercontractility, impaired relaxation and higher intracellular Ca2+ transients that were fully rescued with variant correction.

###### Patient CMs showed altered ATPase activity

*MYH7* variants, especially those located in the head and neck domains, can increase myosin-ATPase activity resulting in increased hydrolysis of myosin-bound ATP to ADP and inorganic phosphate thereby increasing the binding of myosin to actin and increasing contractility.^28; 29^ We assessed ATPase activity in CM lysates by measuring inorganic phosphate generation after adding an ATPase substrate. ATPase activity was higher in 80, 81 and 82 patient CMs (p<0.05 vs. PGPC17) (**Figure 3H**), but not in 83 CMs that harbored the additional *TNNI3* variant. ATPase activity normalized in variant-corrected patient CMs (p<0.05 vs. patient CMs) (**Figure 3H, Table S1)**.

###### Patient CMs showed altered myosin-actinin binding

We performed co-immunoprecipitation to assess the binding affinity of MYH7 to α-actinin in patient CMs compared to controls.

##### MYH7 variant CMs

Baseline α-actinin, MYH7, and MYBPC3 expression was not different in cell lysates from *MYH7* variant patient CMs (80, 81) compared to PGPC17 (**Figure 4A, D, G**). Immunoprecipitation with anti-MYH7 antibody revealed more abundant α-actinin pull-down (p<0.05 vs. PGPC17) and was normal in variant-corrected CMs (p<0.05 vs. patient CMs) (**Figure 4B-C, Table S1**). A similar result was seen when using anti-α-actinin antibody to pull down MYH7 protein, i.e., more abundant MYH7 pull-down in 80 and 81 compared to PGPC17 (**Figure 4E-F, Table S1**).

**Figure 4.**
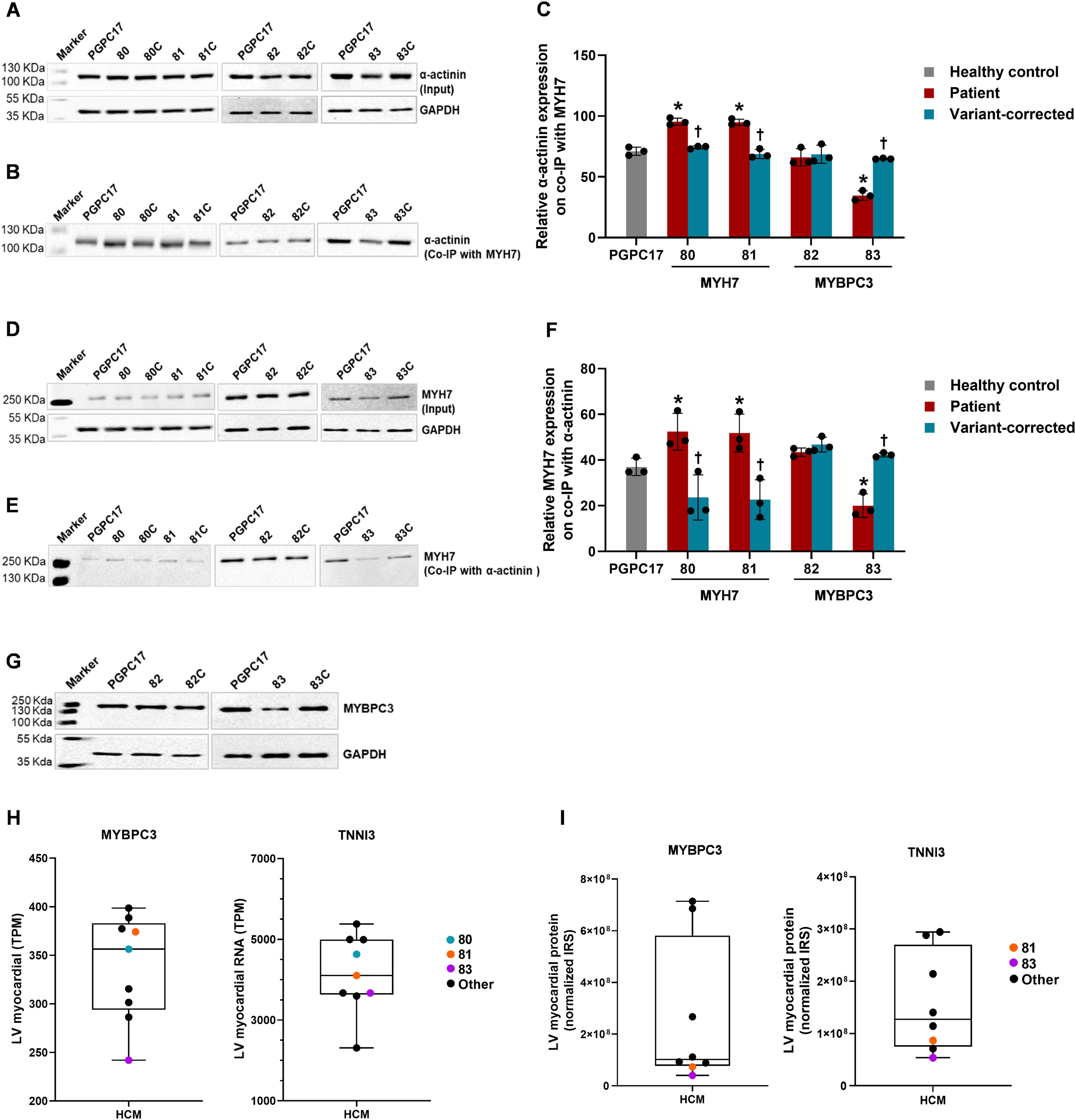
Myosin-actinin co-immunoprecipitation in iPSC-CM lysates. **A. Baseline actinin protein expression** by Western blots was not different in cell lysates from patient compared to control and variant-corrected CMs (representative blots shown). GAPDH was used as the house keeping protein. **B, C. Actinin protein expression on co-immunoprecipitation with anti-MYH7 antibody** (representative blots and quantification relative to GAPDH): There was higher actinin protein expression on co-IP with MYH7 antibody in 80 and 81 patient CMs but not in 82 patient CMs compared to PGPC17 control and variant-corrected CMs. Actinin protein expression was lower on co-IP with MYH7 antibody in 83 patient CMs compared to PGPC17 control and variant-corrected CMs. **D. Baseline MYH7 protein expression** on Western blots was not different in cell lysates from control, patient and variant-corrected CMs. **E, F. MYH7 protein expression on co-immunoprecipitation with anti** α**-actinin antibody:** There was higher MYH7 protein expression on co-IP with actinin antibody in 80 and 81 patient CMs, but not in 82 patient CMs compared to PGPC17 control and variant-corrected lines. MYH7 protein expression was lower on co-IP with actinin antibody in 83 patient CMs compared to PGPC17 control and variant-corrected CMs. **G. Baseline MYBPC3 expression in *MYBPC3* mutant lines:** MYBPC3 protein levels were not different in 82 patient CMs compared to controls but was lower in 83 compared to PGPC17 and variant-corrected CMs. **H-I. MYH7, MYBPC3 and TNNI3 mRNA and protein expression in LV myocardium from HCM patients**. **Boxplots showing the mean and range (minimum to maximum) expression. H.** RNA sequencing performed on LV myocardium from patients 80, 81 and 83 and 6 other HCM patients showed lower *MYBPC3* mRNA levels and borderline low *TNNI3* levels in patient 83 compared to other HCM patients. **I.** Proteomic expression on mass spectrometry on LV myocardium from patients 81, 83 and 6 other HCM patients revealed lowest MYBPC3 and TNNI3 protein expression in 83 compared to other patients. *p<0.05 patient vs. PGPC17, ^†^p<0.05 patient vs. corrected. n=3 independent experiments. Error bars represent standard deviation. 80, *MYH7* V606M; 81, *MYH7* R453C; 82, *MYBPC3* G148R; 83, *MYBPC3* P955fs and *TNNI3* A157V. CM, cardiomyocytes; MYH7, myosin heavy chain 7; Co-IP, co-immunoprecipitation; LV, left ventricular; TPM, transcript per million; IRS, internal reference scaling.

##### MYBPC3 variant CMs

Baseline α-actinin and MYH7 expression was not different in cell lysates from *MYBPC3* variant patient CMs (82, 83) compared to PGPC17 (**Figure 4A, D, G**). Baseline MYBPC3 protein level was also not different in 82 CMs (which harbored a missense variant) but was expectedly lower in 83 CMs (which harbored a frame-shift variant) compared to PGPC17 and variant-corrected controls (**Figure 4G**). Unlike *MYH7* variant CMs, myosin-actinin binding was not different in 82 patient CMs compared to PGPC17. Of note, myosin-actinin binding was 2-fold lower in 83 patient CMs compared to PGPC17 and normalized in variant-corrected CMs (p<0.05). A similar result was seen when using anti-α-actinin antibody to pull down MYH7 protein (**Figure 4E-F, Table S1**).

Therefore, only *MYH7* CMs showed higher myosin-actinin binding in the context of higher ATPase activity while the *MYBPC3* CMs did not show an increase in myosin-actinin binding. In fact, 83 patient CMs with both *MYBPC3 and TNNI3* variants showed lower myosin-actinin binding.

#### Protein expression in left ventricular myocardium of double-variant patient

To better understand the differences in baseline phenotype in 83 patient CMs with *MYBPC3 and TNNI3* variants, we analyzed MYBPC3 and TNNI3 gene and protein expression in patient left ventricular myocardium using RNA sequencing and mass spectrometry. We compared expression in 83 vs. 80, 81 (82 not available) and 6 other HCM pediatric patients from our biobank without *MYBPC3* variants. *MYBPC3* mRNA levels were the lowest in patient 83 compared to other HCM patients. *TNNI3* mRNA expression was also low in 83 (**Figure 4H**). This was confirmed with mass spectrometry which revealed lowest MYBPC3 and TNNI3 protein expression in 83 compared to the other patients (**Figure 4I**). These findings derived directly from patient myocardium suggest that reduced MYBPC3 and TNNI3 expression may have influenced the functional phenotype of 83 patient CMs.

In summary, we successfully modeled childhood onset HCM using patient iPSC-derived CMs, identified functional differences by genotype, and confirmed that the abnormalities were related to the pathogenic variants using variant-corrected isogenic CMs.

#### Effect of standard and myosin-targeted drugs on patient CMs

We compared the ability of a targeted myosin ATPase inhibitor, MYK-461, and of non-targeted drugs, i.e., verapamil, a Ca2+ channel blocker, and metoprolol, a beta-blocker, to rescue functional abnormalities in patient CMs. We examined the acute effects on CM contractility, relaxation, Ca2+ transients and ATPase activity using 3 doses for MYK-461 and the highest tolerated dose that had no cytotoxic effects for verapamil and metoprolol (**Figure 5A**). Drug response was assessed by comparing effects at baseline, i.e., pre-treatment, and 3 hours after drug administration, the time point at which maximal drug effects were seen.

**Figure 5.**
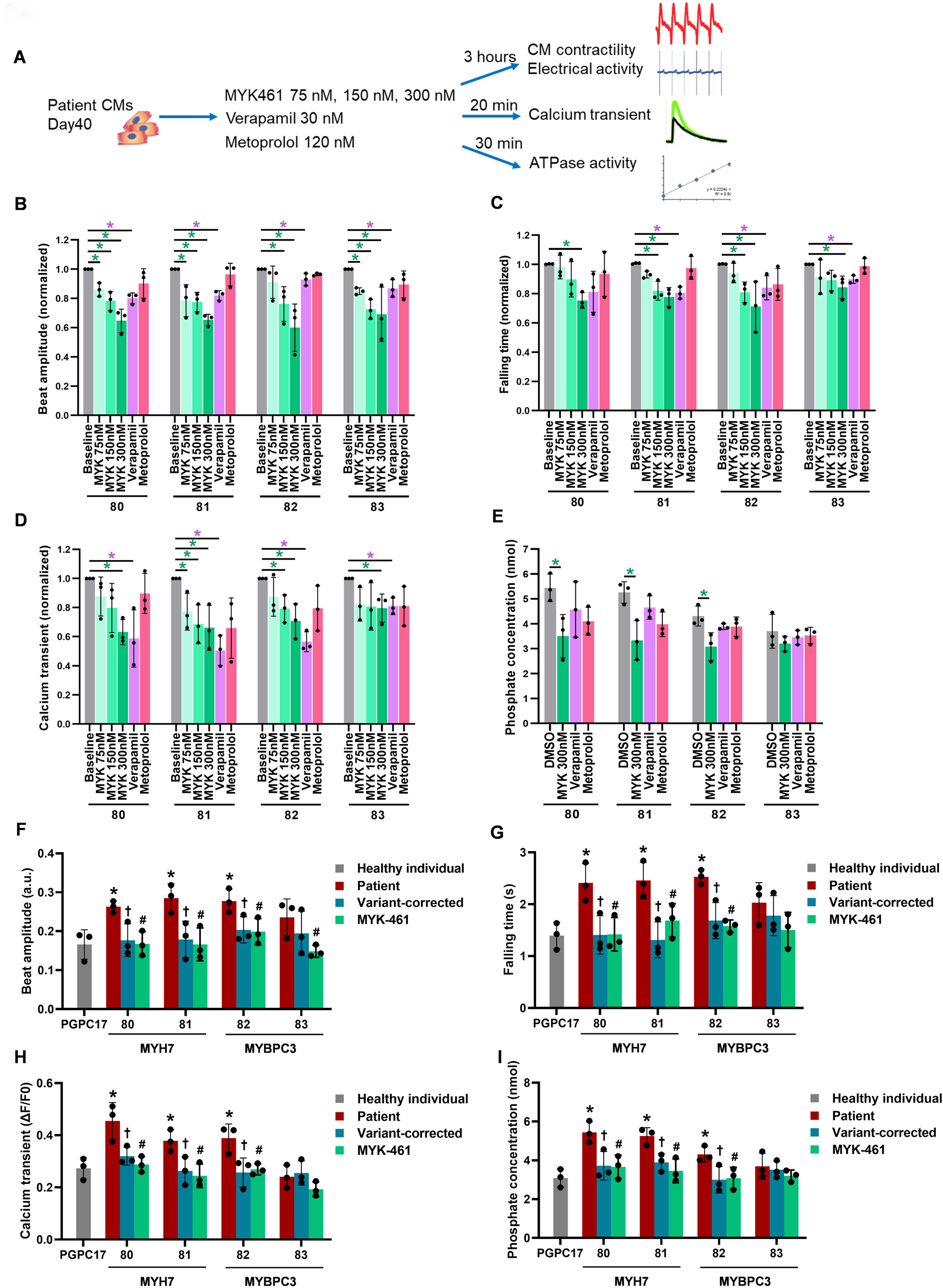
Drug responses to MYK-461, verapamil and metoprolol in HCM patient iPSC-CMs. **A.** Schematic illustration of drug treatment (dose and duration) and functional assays to assess drug response. **B. Beat amplitude**: MYK-461 caused a dose-dependent decrease in beat amplitude in all HCM patient lines. Verapamil also reduced beat amplitude in all patient CMs but this effect was lower than that seen with highest dose of MYK-461. Metoprolol had no effect. **C. Falling time**: MYK-461 caused a dose-dependent decrease in falling time in all HCM patient CMs. Verapamil decreased falling time in 81, 82, and 83 CMs but not in 81, while metoprolol had no effect on falling time. **D. Calcium transients:** MYK-461 caused a dose-dependent decrease in calcium transients in all four patient CMs but the response was blunted in 83 CMs. Verapamil decreased calcium transients in all 4 patient CMs but the response was blunted in 83 CMs. Metoprolol had no effect. **E. ATPase activity**: MYK-461 decreased ATPase activity in 80, 81 and 82 CMs but not in 83 CMs. Verapamil and metoprolol had no effect. **F-I. Effect of highest MYK-461 dose compared to PGPC17 control and to variant-corrected lines. F. Beat amplitude:** MYK-461 (300 nM) decreased abnormal beat amplitude to the same level as PGPC17 and variant-corrected lines in all HCM patient CMs. **G. Falling time, H. Calcium transients,** and **I. ATPase activity** were decreased by MYK-461 treatment in 80, 81 and 82 CMs to levels seen in PGPC17 and variant-corrected CMs. *p<0.05, ^†^p<0.05 patient vs. corrected, ^#^p<0.05 patient vs. MYK-461. n=3 independent experiments, using 4 technical replicates for each experiment. Error bars represent standard deviation. **B-D:** Drug treatment values were normalized to pre-treatment. 80, *MYH7* V606M; 81, *MYH7* R453C; 82, *MYBPC3* G148R; 83, *MYBPC3* P955fs and *TNNI3* A157V. CM, cardiomyocytes; a.u., arbitrary unit; ΔF/F0, ratio of peak fluorescence intensity to baseline intensity.

##### Drug effects on contractility and relaxation

MYK-461 caused a dose-dependent decrease in beat amplitude in all 4 patient CMs (60-70% at highest dose; p<0.05 vs. pre-treatment) (**Figure 5B**). Verapamil also reduced beat amplitude in all 4 patient CMs (p<0.05 vs. pre-treatment) but the effect was lower with reduction to 80-90% of baseline compared to the highest dose of MYK-461 (p<0.05 MYK 300nM vs. verapamil) (**Table S2**). Metoprolol did not lower beat amplitude in any patient CMs (**Figure 5B**). MYK-461 lowered falling time in all 4 HCM patient CMs at the highest dose (p<0.05 vs. pre-treatment). Verapamil also reduced relaxation time in 80, 81 and 82 patient CMs (p<0.05 vs. pre-treatment) but not in 83 patient CMs (which did not have relaxation abnormalities at baseline). Metoprolol did not affect relaxation time (**Figure 5C**). Beat rate, FPA and FPD were not altered by the drugs in any patient CMs. We further compared the effect of the highest dose of MYK-461 to PGPC17 and to variant-corrected controls. MYK-461 decreased abnormal beat amplitude to the same level as PGPC17 and variant-corrected controls in all HCM patient CMs (**Figure 5F**). Relaxation time was normalized by MYK-461 in 80, 81 and 82 CMs to levels seen in PGPC17 and variant-corrected controls (p>0.05 vs PGPC17 and vs. corrected CMs) (**Figure 5G**).

##### Drug effects on Ca2+ transients and ATPase activity

MYK-461 and verapamil reduced Ca2+ transients in all 4 patient CMs (p<0.05 vs. pre-treatment) (**Figure 5D**). MYK-461 reduced inorganic phosphate generation, i.e., marker of ATPase activity in 80, 81, and 82 patient CMs (p<0.05 vs. pre-treatment) but not 83 patient CMs which had normal ATPase activity at baseline. As expected, verapamil and metoprolol did not affect ATPase activity (**Figure 5E**). The reduction in Ca2+ transients and ATPase activity with high dose of MYK-461 was comparable to that seen in PGPC17 and variant-corrected CMs (**Figure 5H-I**).

Absolute values for all drug responses are shown in **Table S2**.

### Phenotype and drug response in engineered cardiac Biowires from patient and variant-corrected CMs

We evaluated the HCM phenotype and the effect of MYK-461 using patient iPSC-derived cardiac tissues (Biowires) to investigate the effects on contractile force which is not possible in monolayer cultures. This was done on one *MYH7* variant (80) and one *MYBPC3* variant (82) patient and corrected CMs that exhibited a significant phenotype in cell culture. Biowires were generated by combining CMs and cardiac fibroblasts (10:3 ratio) with hydrogel within microwells as previously described.^30^ The Biowires underwent remodeling and compaction over 7 days to form cylindrical trabecular strips (**Figure 6A**). Compaction rate from day 2 to day 7 was not different between patient and variant-corrected Biowires (**Figure 6B**). Immunostaining was used to confirm the expression of sarcomeric α-actinin and MLC2v (**Figure 6C**). Nuclear area, a surrogate for cell size,^31^ was larger in both 80 and 82 patient Biowires compared to variant-corrected Biowires (p<0.05) (**Figure 6D**). Fiber length was shorter in patient compared to variant-corrected Biowires (p<0.05) indicating less organized sarcomere protein arrangement in patient CMs (**Figure 6E**).

**Figure 6.**
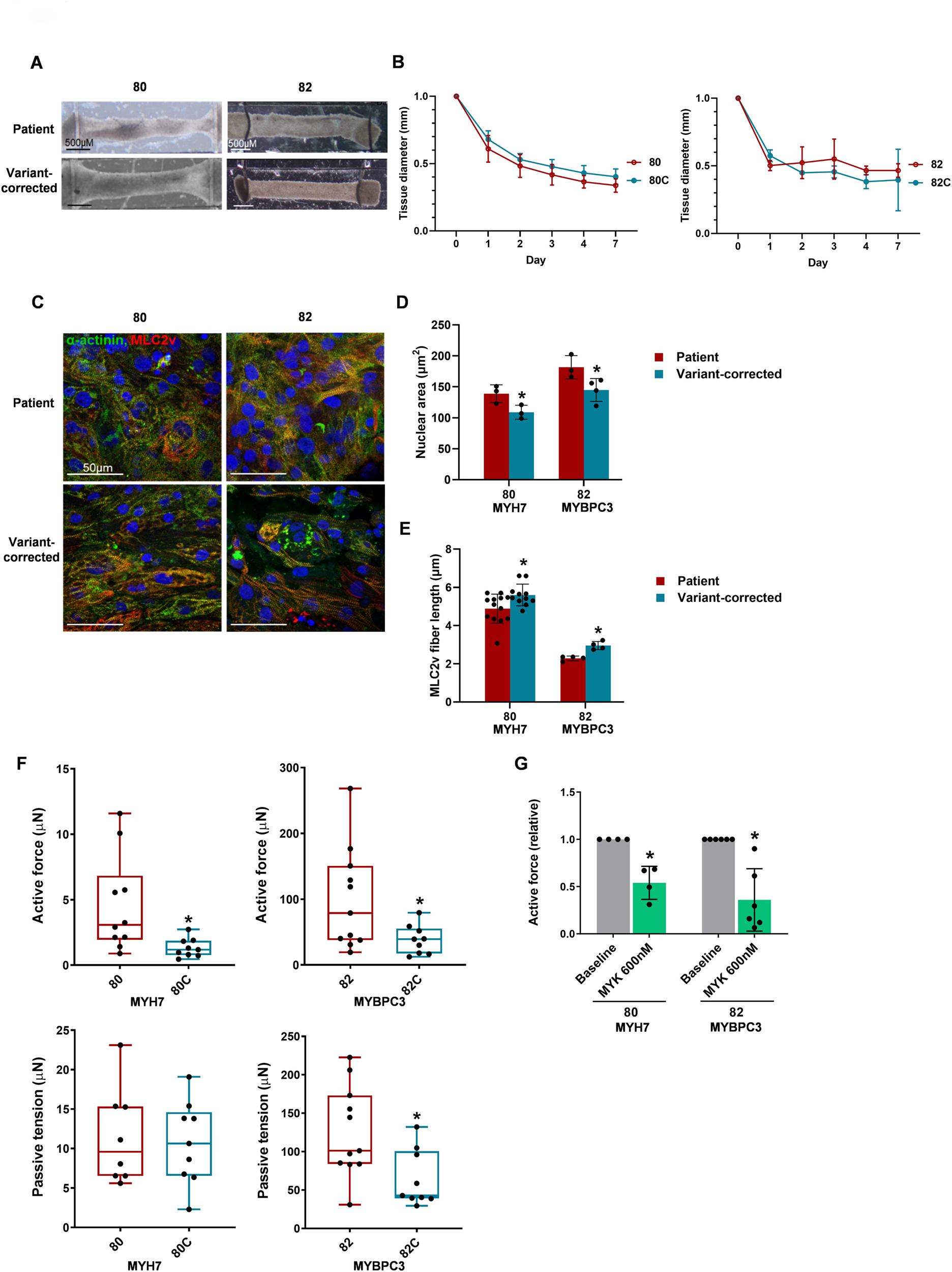
Cardiac Biowires from patient and variant-corrected iPSC-CMs. **A. Bright field images** of Biowires (day 7 of seeding) from 80 and 82 patient CMs within microwells. **B. Tissue compaction** increased from day 0 to day 7 in patient and variant-corrected Biowires. **C. Confocal images** (representative) of Biowire tissues immunostained for α-actinin and MLC2v. **D. Nuclear area** was larger in patient Biowires compared to variant-corrected Biowires. **E. Fiber length** (MLC2v) was lower in patient Biowires (suggesting sarcomere disorganization) compared to variant-corrected Biowires. **F. Active force** was higher in patient Biowires compared to variant-corrected Biowires. Passive tension was only higher in 82 patient Biowires compared to variant-corrected Biowires. **G. MYK-461** decreased active force in both patient Biowires relative to untreated Biowires. *p<0.05. n=3 to 13 biological replicates. Error bars represent standard deviation. 80, *MYH7* V606M; 82, *MYBPC3* G148R. CMs, cardiomyocytes; MLC2v, ventricular myosin light chain 2.

Contraction dynamics of engineered Biowires were assessed under electrical stimulation. The electrical excitation threshold and maximum capture rate were not different between patient and variant-corrected Biowires, consistent with findings in CM monolayers (**Figure S6A-B**). Active force was 2.6 to 3.4-fold higher in patient compared to variant-corrected Biowires (p<0.05) (**Figure 6F**). Passive tension was lower only in 82 patient compared to corrected Biowires (p=0.01) (**Figure 6F**). Absolute values are provided in **Table S1**.

We studied the effect of MYK-461 using this tissue model. Since the outer layer of Biowires is fibroblast-rich, drug diffusion into tissues can be slower than in CM cultures. Hence we used a higher dose of 600 nM of MYK-461 for 24 hours. MYK-461 decreased active force by in patient Biowires (p<0.05 vs pre-treatment) (**Figure 6G**). There was no effect of the drug on excitation threshold or maximum capture rate (**Figure S6A-B**). Absolute values are provided in **Table S2**.

In summary, MYK-461 showed complete rescue of abnormal contractility, relaxation, Ca2+ transients, and ATPase activity in *MYH7* and *MYBPC3* variant CMs in culture, and also in engineered tissues. Verapamil improved contractility and relaxation abnormalities although the response was partial compared to MYK-461, and did not alter ATPase activity. Metoprolol had no effect. The targeted myosin ATPase inhibitor therefore showed more consistent and complete rescue of the abnormal phenotype compared to non-targeted drugs in iPSC-CMs from pediatric HCM patients.

## DISCUSSION

We demonstrated morphological and functional hallmarks of HCM in iPSC-derived CMs and cardiac Biowire tissues from patients harboring disease-causing pathogenic variants in two of the most commonly mutated genes – *MYH7* (V606M, R453C) and *MYBPC3* (G148R, P955fs). Importantly, we corrected all four variants to generate isogenic CMs that showed complete rescue of the abnormal phenotype. This enabled us to confirm that the phenotype was indeed caused by the genetic variant and also allowed us to compare pharmacological rescue to genetic rescue. The main finding of the study is that drug responses in pediatric HCM patient CMs varied by genotype and by drug class, and that a myosin ATPase inhibitor showed complete and consistent rescue of the HCM phenotype compared to non-targeted drugs highlighting the need for trials of this drug in a pediatric population.

The use of a platform that allowed serial assessment of both mechanical and electrophysiological abnormalities and the ability to correlate with abnormalities in Ca2+ signaling and ATPase activity are a strength of this study. The morphological phenotype of myocyte hypertrophy and sarcomere disorganization were seen in all patients, but hypercontractility, impaired relaxation, and altered Ca2+ handling were only seen in patients with *MYH7* (V606M, R453C) and *MYBPC3* (G148R) single variants. Several studies have reported morphological and functional abnormalities in HCM patient iPSC models harboring *MYH7* or *MYBPC3*variants.^10; 11; 16–18; 20; 21^ *MYH7* R543C and V606M have been modeled in gene edited control iPSCs,^9; 12–14^ but have not been studied in patient iPSCs, while *MYBPC3* G148R and P955fs variants have not been studied in gene-edited or patient iPSCs. Therefore our study provides insights into genotype-phenotype association through the use of patient derived iPSCs. Further, only one previous report performed gene editing^21^ to generate isogenic controls which is critical to attribute the observed phenotype to the disease-associated variant.

Despite similar HCM morphological phenotype, there were differences in CM function between genotypes likely related to differences in genotype. Specifically, only patients with *MYH7* variants (80, 81) showed both increased ATPase activity and increased myosin-actinin binding, both of which were rescued with variant correction. Both the *MYH7* variants were located in the head domain close to the ATPase and the actin binding sites on myosin which may explain the observed increase in ATPase activity and myosin-actinin binding in these patient iPSC-CMs. Unlike the patients with *MYH7* variants, patient 82 with the *MYBPC3* G148R single variant showed a modestly higher ATPase activity while patient 83 showed no increase in ATPase activity. Neither of the patients with *MYBPC3* variants demonstrated increased myosin-actinin binding suggesting that the mechanism of hypercontractility in *MYBPC3* variant positive patients was independent of myosin-actinin binding.

Patient 83 who harbored two pathogenic variants, *MYBPC3* P955fs and *cTNNI3* A157V, showed a phenotype that was distinct from the other 3 patients. Patient 83 CMs showed morphological abnormalities but no hypercontractility or impaired relaxation, no increase in calcium transients, no increase in ATPase activity and importantly, showed lower myosin-actinin binding. We speculate that the reduced MYH7-actinin binding may be related to reduction in MYBPC3 protein (seen both in patient CMs and in patient myocardium) rather than to the effects of the co-existing *TNNI3* variant since the lower myosin-actinin binding was rescued by correction of the *MYBPC3* variant alone. Nonetheless, it is not possible to completely exclude a confounding effect of the *TNNI3* variant on baseline phenotype. Cardiac troponin I is the inhibitory subunit of the thin filament regulatory protein troponin that regulates cardiac muscle contraction in response to Ca2+.^32^ *TNNI3* A157V has been reported in over 20 individuals with HCM,^33^ as well as in an individual with dilated cardiomyopathy,^34^ and has been reported to cause a decrease in TNNI3 protein level (similar to our patient), but its effect on CM function has not been previously studied.^35^ Overall, our findings suggest that even in patients with HCM caused by sarcomere gene variants, the pathophysiologic mechanisms that contribute to the disease phenotype likely differ. This could have implications for the response to different pharmacological agents.

In that regard, we found that a targeted small molecule myosin-ATPase inhibitor, MYK-461, was effective at rescuing the in vitro morphological and functional abnormalities in 3 of the 4 patient CMs (80, 81, 82) that had baseline abnormalities in CM contraction, relaxation, Ca2+ signaling, and ATPase activity. In fact, it was more effective than a standard calcium channel blocker, verapamil, or a beta-blocker, metoprolol, in rescuing the HCM phenotype in patient CMs. MYK-461 treatment resulted in return of functional parameters to normal i.e., to that seen in variant-corrected and PGPC17 healthy control CMs. Like MYK-461, verapamil was able to rescue hypercontractility, impaired relaxation, and Ca2+ transients in affected patient CMs but the effect, at the highest tolerated dose, was lower than that seen with MYK-461. Also verapamil did not alter ATPase activity i.e., its effects on contractility were independent of ATPase activity. This may explain in part the lower efficacy of verapamil compared to MYK-461. Metoprolol, which did not affect calcium transients or ATPase activity, did not rescue contractile or relaxation abnormalities. This is consistent with a previous study that showed verapamil rescued Ca2+-handling abnormalities and arrhythmia in *MYH7* R663H pediatric patient CMs, while metoprolol did not.^10^ In general, although some previous studies have reported the effects of MYK-461 in iPSC-CMs or engineered cardiac tissues, they have been primarily from adult patients and direct comparison with non-targeted drugs and with isogenic controls has not been reported.^9, 36^

Since this was a study of acute drug effects, we did not assess if longer drug treatment would rescue the morphological phenotype of CM hypertrophy and sarcomere disorganization. Nonetheless, the acute findings suggest that MYK-461 and, to a lesser extent, verapamil, can rescue CM functional and biochemical abnormalities in pediatric patient CMs but this effect is influenced by the genotype. The efficacy was highest in those with missense variants in *MYH7* or *MYBPC3* and was not seen in the patient with *MYBPC3* haploinsufficiency.

A study limitation is that only iPSCs from male patients were analyzed. A majority (70%) of affected HCM patients are males. We were limited in this study to families that consented to the biobank with *MYH7* or *MYBPC3* variants who provided a DNA sample for sequencing and a blood or skin sample for reprogramming. None of the female patients in our biobank met the above criteria. Therefore we were unable to analyze sex-related differences in CM phenotype.

In summary, we demonstrated the utility of patient and variant-corrected patient iPSC-CMs for modeling HCM and testing new therapeutic compounds and validated our findings in bioengineered cardiac tissues. Our findings have important clinical implications. First, our study demonstrated the *in vitro* efficacy of a myosin ATPase inhibitor in patients with childhood-onset HCM, and highlighted the need for clinical trials in of myosin-targeted drugs in children.^8; 37; 38^ Further, since the response to drug therapy appears to be influenced by the underlying genotype, clinical drug-gene interaction studies are indicated in order to guide genotype-targeted uptake of these new precision therapies.

## ACKNOWLEDGEMENTS

We acknowledge the Centre for Commercialization of Regenerative Medicine in Toronto for iPSC reprogramming and gene editing and the Centre for Applied Genomics at the Hospital for Sick Children for performing RNA sequencing. We thank Leanne Wybenga-Groot and Michael Moran from Sickkids Proteomics, Analytics, Robotics and Chemical Biology Centre at the Hospital for Sick Children for conducting mass spectrometry. We thank patient participants in the Heart Centre Biobank Registry for contributing data and samples. We thank Joseph Wu, Stanford Cardiovascular Institute, for sharing human iPSC lines (SCVI 4, SCVI 480) funded by NIH R24 HL117756 that were used by us to optimize our iPSC-CM differentiation protocols and assays.

## AUTHOR CONTRIBUTIONS

Conceptualization, J.E. and S.M.; methodology and investigation, C.K., A.S., G.M., Y.Z., E.Y.W., N.P., W.W., M.R., J.E. and S.M.; writing original draft, C.K.; writing and editing, C.K., A.S., G.M., E.Y.W., C.A.S., M.R., J.E. and S.M.; funding acquisition, F.B., C.A.S., M.R., J.E. and S.M.; supervision, M.R., J.E. and S.M. All authors reviewed and revised the final version of the manuscript.

## INCLUSION AND DIVERSITY

We support inclusive, diverse, and equitable conduct of research.

## METHODS

### Resource availability

#### Lead contact

Further information and requests for resources should be directed to the lead contact, Dr. Seema Mital.

### Materials availability

All unique reagents generated in this study are available from the lead contact with a completed materials transfer agreement.

### Data and code availability

Sequencing data are deposited in the European Genome-Phenome Archive (EGA) under accession EGAS00001004929, and are available for download upon approval by the Data Access Committee. Additional data generated or analyzed during this study are included in the supplementary information files, and additional raw data used for figures and results are available from the corresponding author on reasonable request. All computational tools used in this study are available for download as commercial or open-source software and are detailed in Methods. Detailed parameters of all functions used for each tool are available in the Methods.

### Experimental Methods

#### Generation of patient and CRISPR-Cas9 gene-corrected iPSCs

Skin fibroblasts or lymphoblasts were obtained from 4 pediatric patients with HCM recruited through the SickKids Heart Centre Biobank Registry (Toronto, ON, Canada). Studies were approved by the Hospital for Sick Children institutional review board, and written informed consent was obtained from all patients and/or their parents/legal guardians.^24; 25^ Patients harbored the following myosin variants - *MYH7* p.V606M (c.1816G>A) (80), *MYH7* p.R453C (c.1357C>T) (81), *MYBPC3* p.G148R (c.G442A) (82), and *MYBPC3* p.P955fs (c.2864_2865del) (83). Variants were identified on clinical testing as well as on research whole genome sequencing and met the American College of Medical Genetics criteria for pathogenicity.^25^ These somatic cells were reprogrammed at the Centre for Commercialization of Regenerative Medicine (CCRM) (Toronto, Canada) into human induced pluripotent stem cells (iPSCs) using a Sendai virus vector carrying transcription factors Oct4, Sox2, Kif4, and c-Myc.^26^ To generate isogenic controls, patient iPSCs were gene edited at CCRM using CRISPR/Cas9 mediated Homologous Recombination. In brief, a Cas9 guide RNA (gRNA) close to the targeted mutation and a 100 base single stranded oligo DNA repair template containing the gene correction were designed. Cells were co-transfected with Cas9-gRNA RNP complex together with the repair oligo and Puromyin-GFP plasmid^39^ (Addgene, Watertown, MA) using Lipofectamine Stem transfection reagent (ThermoFisher, Waltham, MA). Transfected cells were subjected to Puromycin selection for 48 hours to enrich co-transfected cells. Single cell colonies were then screened for gene correction using a ddPCR assay using probes corresponding to the wild-type or mutant sequence. Variant correction was further confirmed by PCR and Sanger sequencing of the region of interest. To rule out off target cleavage by the guide RNA, PCR and Sanger sequencing was performed for the top 5 potential off-target sites for the corresponding gRNAs. No off-target activity was identified in any of the gene-corrected clones. Mycoplasma was tested using the MycoAlert detection kit (Lonza, Rockland, ME). Cell identity was confirmed using short tandem repeats by PCR profiling. Karyotyping was conducted using G-banding analysis to confirm chromosomal integrity. Cells were tested for pluripotency using flow cytometry for expression of SSEA4, Tra-1-60, OCT4, and SOX2. We used previously published iPSCs (PGPC17) as healthy controls.^27^ Whole genome sequencing of the PGPC17 donor showed this genome to be devoid of any pathogenic variants in cardiac-disorder related genes^40^ (**Figures S2-5)**.

#### Cardiomyocyte differentiation

Patient and control iPSCs were differentiated into CMs in accordance with an established differentiation protocol using STEMdiff Cardiomyocyte Differentiation Kit^27^ (STEMCELL Technologies, Vancouver, BC). The CMs were then maintained from day 8 using a combination of Stem Cell Technologies and iCell Maintenance media (FUJIFILM Cellular Dymamics, Madison, WI) where Stem Cell Technologies medium was gradually decreased by 25% and iCell medium increased by 25% every other day until CMs were maintained in 100% iCell medium (day 14-16). iCell medium can effectively remove non-cardiomyocytes in differentiated cells. On day 16, cells were dissociated and reseeded in StemCell Technologies cardiomyocyte support medium to improve their attachment. 48 hours later, cells were fed and maintained in iCell Maintenance medium with a change of media every 48 hours. Cellular assays were performed at day 38-42, when cells exhibited a more mature phenotype involving regular contractility and stable electrical signals. All experiments were done using 3 biological replicates (3 independent CM differentiations). Data was plotted from the average of 4 technical replicates for each of 3 biological replicates.

#### Immunofluorescence staining

CMs were fixed with 4% paraformaldehyde at day 38, permeabilized with 1% Triton X and blocked with 5% bovine serum albumin. Immunostaining was conducted with mouse α-actinin and rabbit myosin light chain 2, MLC2v antibodies (1:500, Abcam, Cambridge, MA). CMs were then incubated with Alexa Fluor 488 and Alexa Fluor 555 conjugated antibodies (1:1000, ThermoFisher). DAPI was used as counterstaining for nucleus. Images were taken from 8 fields per well in 4 wells at 2 different magnifications for each biological replicate using a spinning disk confocal microscope (Quorum Technologies, Guelph, ON) and the Volocity software (PerkinElmer, Waltham, MA). Cell size was measured using ImageJ (n≈600 cells/biological replicate per cell line) (National Institutes of Health, Bethesda, MD). Sarcomere organization was assessed by line-scan analysis of MLC2v and α-actinin alignment (ImageJ) (n≈150 cells/biological replicate per cell line).

#### Contractility, relaxation and electrophysiological assessment

The xCELLigence real-time cell analysis (RTCA) CardioECR system (Agilent, Santa Clara, CA) was used to measure beat rate, contractility and relaxation. 48 well xCELLigence E-plates were coated with fibronectin for one hour. Before adding cells, a baseline recording was taken while only media was present in the well. 4×10^4^ dissociated CMs were seeded in each well on day 16. Cells were maintained in iCell Maintenance medium, with a change of media every 48 hours. From day 30 to 40, continuous 20 second recordings were obtained every 3 hours to assess baseline phenotype. On day 40, 20 second recordings were acquired every 10 minutes for 3 hours to measure CM contractility, relaxation and electrical activity. The platform uses a plate coated with interdigitated impedance microelectrodes that captures spontaneous contraction based on the change in mechanical impedance of CMs. Beat rate (beats per minute) is the total number of positive peaks in one minute. Beat amplitude (arbitrary unit) is the measure from each negative peak to the following positive peak. Falling time (ms) is the time it takes for the cell index to decrease from 80% to 20% amplitude in the downstroke stage of each beat. The plate also contains two field potential electrodes that capture electrical activity, field potential amplitude (FPA) and field potential duration (FPD). FPA (mV) is the amplitude between field potential (FP) spike negative peak and FP spike positive peak. FPD (ms) is the duration between FP spike negative peak to the end of the FP wave.

#### Calcium imaging

4×10^4^ cells per well were seeded at day 16 in gelatin-coated 96-well plates. At day 40, CMs were loaded with 1 µM Fluo-4 AM (Life Technologies, Carlsbad, CA) in HBSS solution for 30 minutes at 37°C and washed. For drug screens, cells were treated with drugs and DMSO only (as negative control) for 20 minutes. Live imaging was performed with a Nikon TE2000 microscope at 37°C using Volocity software (Nikon Corporation, Tokyo, Japan; PerkinElmer). Images were captured with 488 nm excitation and 516 nm emission at a rate of 20 images per second for 30 seconds. Image stacks were converted to videos with Volocity and regions of interest (4 per technical replicates) were analyzed for changes in fluorescence intensity (ΔF/F0) with the resting fluorescence value F0 set as the minimal value recorded.

#### ATPase assay

1×10^6^ cells were seeded per well in gelatin-coated 6 well plates. Cells were lysed and homogenized at day 42 with ATPase assay buffer. ATPase activity was assessed using a colorimetric ATPase assay kit (Abcam) by measuring inorganic phosphate generation in CMs. An ATPase substrate was added hydrolyzing ATP, thereby releasing ADP and a free phosphate ion. Linked reactions enable a stable chromophore to be generated at an optical density of 650 nm, which is proportional to the amount of enzyme activity present.

#### MYH7 and α-actinin co-immunoprecipitation assay

1×10^6^ cells were seeded per well in gelatin-coated 6 well plates. CMs were lysed at day 40 in RIPA buffer (1 mM Tris, pH 7.4, 1% Triton X-100, 5 M sodium chloride, 0.25% sodium deoxycholate, 0.5 mM EDTA, 0.1% SDS) and a cOmplete™, Mini, EDTA-free Protease Inhibitor Cocktail (Roche, Mississauga, ON). After centrifugation at 3000g for 3 min, 5% of supernatant was used as input for baseline protein expression. To pull down target proteins, supernatants were incubated with either α-actinin antibody (1:500, Abcam) or anti-MYH7 antibody (1:500, R&D systems, Minneapolis, MN) by shaking gently overnight at 4 °C. Immune complexes were precipitated using either protein A or G agarose beads (Cell Signaling Technology, Danvers, MA) for 4 h at 4 °C. Immune complexes were pelleted, washed 3 times in washing buffer (25 mM Tris, 1 mM EDTA, 0.15 M NaCl, 5% glycerol; pH 7.4, 1% NP-40), 3 times in 1x-PBS and then heated at 65 °C for 5 min in loading buffer. Associated protein complexes were eluted from the agarose beads. Reactive bands were visualized using Pierce™ ECL Western Blotting Substrate (ThermoFisher) after SDS–PAGE and immunoblotting with the reciprocal antibodies.

#### Myocardial gene and protein expression

*RNA sequencing* was performed in left ventricular myocardial samples available from 9 HCM patients in our biobank to measure *MYBPC3* and *TNNI3* expression. Total RNA was extracted from myocardial samples using the RNeasy Mini kit (QIAGEN, Toronto, Canada). RNA sequencing was performed using Illumina HiSeq 2500 platform through The Centre for Applied Genomics, Hospital for Sick Children, Toronto. The RNA sequencing expression samples were processed using an in-house pipeline previously described.^41^ Differential expression between 3 HCM samples with *MYH7* and *MYBPC3* variants of interest and 6 other HCM samples without these variants was detected using DESeq2 BioConductor package.^42^ In subsequent analyses gene expression was represented using transcript per million (TPM) values. Boxes mark the 25th percentile (bottom), median (central bar) and 75th percentile (top). Whiskers represent extreme values.

##### Mass spectrometry

Mass spectrometry on left ventricular tissue lysates from 8 HCM patients was performed in the SickKids Proteomics, Analytics Robotics & Chemical Biology Centre as previously described.^43^ Briefly, proteins were extracted by chloroform-methanol precipitation and digested with 2 μg trypsin/Lys-C mixture (Promega, Madison, WI). Samples were labeled with 10-plex tandem mass tag reagents (ThermoFisher Scientific) and separated into 60 fractions using high-pressure liquid chromatography. Fractions were dissolved in 0.1% formic acid and loaded into Evotip C18 trap column (Evosep Biosystems, Odense, Denmark). Peptides were analyzed using Orbitrap Fusion Lumos Tribrid Mass Spectrometer (ThermoFisher Scientific). Automatic gain control targets were 4×10^5^ with a maximum ion injection time of 50 ms. Data was acquired using MultiNotch synchronous precursor selection MS3 scanning for tandem mass tags. Total proteome raw data was analyzed as individual batches using scripts and methods from Phillip Wilmarth’s GitHub PAW-pipeline repository (github.com/pwilmart/PAW_pipeline). Data was searched against Uniprot Human proteome (UP000005640, Oct 2021, 20600 entries) using Comet search engine (v2016.01). Search results were filtered using a false discovery rate (FDR) of 0.01. The proteomic dataset of 2 HCM samples with *MYH7* and *MYBPC3* variants and 6 other HCM samples was normalized using internal reference scaling (IRS).^44^ Replicate controls were used in each mass spectrometry plex to ensure the absence of batch effects.

#### Assessment of drug responses in CM cultures

At day 40, patient CMs were treated with a myosin ATPase inhibitor, MYK-461 (MedChemExpress, Monmouth, NJ), a Ca2+-channel blocker, verapamil (Sigma-Aldrich, Saint Louis, MO), a beta-blocker, metoprolol (Sigma-Aldrich) or vehicle DMSO as negative control (Sigma-Aldrich). PGPC17 and variant-corrected CMs were included as controls. Three doses were tested for each of the 3 drugs on patient CMs to identify the optimal doses for testing. Three doses that showed a graded effect on beat amplitude without inducing toxicity were used for MYK-461 to assess dose-dependent responses (75 nM, 150 nM and 300 nM). Only the highest tolerated dose that did not result in cytotoxicity was selected for verapamil and metoprolol (50 nM and 150 nM respectfully) to conduct contractility, relaxation, electrophysiology and calcium transient assays. xCELLigence measurements were taken every 10 minutes for 3 hours before and after drug treatment. There was no further drug effect beyond 3 hours. Values derived from the average of recordings over 3 hours pre and post treatment were compared. Calcium transients were recorded after 20 minutes of drug addition and ATPase activity was measured after a 30-minute incubation with 300 nM MYK-461.

#### Biowire generation

To create Biowires, CMs and fibroblasts were co-cultured at a 10:3 ratio for the formation of contractile tissue. Patient and variant-corrected iPSCs i.e., 80, 80C, 82 and 82C were differentiated into CMs using monolayer differentiation protocols.^45^ Dissociated cells were initially plated in 75 cm^2^ flask for 1 hour prior to collection. The attached cells were detached and used as isogenic fibroblast like cells with 0.05% trypsin/EDTA (Life technology). The non-attaching cells were collected and plated on matrigel coated 6-well plates (1:100 dilution) for CM expansion according to the previous reported method.^46^ Briefly, CMs were plated at 40-50,000/cm^2^ and expanded with 2 µM of CHIR in RPMI supplemented with B27. CMs were collected with collagenase II and mixed with previously collected isogenic fibroblasts. The mixed cells were incorporated in collagen hydrogel (5.5×10^7^ cells/mL). Collagen hydrogel was formed by mixing rat tail collagen (153 µL at 9.82 mg/mL, Corning), 1X M199 (50 µL, Sigma-Aldrich), Matrigel (75 µL, BD Biosciences, San Jose, CA), deionized H2O (167 µL), NaOH (5 µL at 10 mM, Sigma-Aldrich) and NaHCO3 (50 µL at 2.3 mM, Sigma-Aldrich). The cell gel mixture was seeded in the microwells of the polystyrenes stripes and was incubated at 37°C and 5% CO2 for 10 mins to allow the hydrogel gelation. Induction 3 Medium (StemPro-34 complete media, 20 mM HEPES, 1% GlutaMAX, 1% Penicillin-Streptomycin (Life Technologies); 213 μg/mL 2-phosphate Ascorbic Acid (Sigma-Aldrich)) was added and changed semi-weekly. Tissues were cultured for 7 days for compaction to occur. Morphology was observed daily from day 0 (seeding day) to day 7 using an Olympus CKX41 inverted microscope. Biowires were cultured for an additional 7 to 14 days prior to doing assays.

#### Biowire immunostaining

Immunofluorescent staining was performed to assess nuclear area as a marker of cell size,^31^ and fiber length as a measure of sarcomere organization. Tissues were fixed with 4% paraformaldehyde, permeabilized with 1% Triton X, and blocked with 5% bovine serum albumin. Immunostaining was performed using mouse anti-α-actinin (1:200, ThermoFisher), rabbit anti-MLC2v (1:200, Abcam) and secondary antibodies Alexa Fluor 488 and Alexa Fluor 647 (1:400, Abcam). DAPI was used as counterstaining for nucleus. Confocal fluorescent microscopy images were generated using an Olympus FluoView 1000 laser scanning confocal microscope (Olympus Corporation, Center Valley, PA).

#### Biowire force analysis

Biowire strips contain a pair of elastic polymer sensors to monitor the contraction dynamics of engineered tissues. Videos of sensors were recorded while Biowires were paced at 1 Hz. Force analysis was performed with MATLAB codes as previously reported.^30^ The outer layer of Biowires is fibroblast-rich, drug diffusion into tissues can be slower than in CM cultures. A higher dose (600 nM) of MYK-461 at 3 and 24 hour timepoints were therefore tested. 24 hour treatment was then selected as the time to capture the optimum response.

### Statistical Analysis

Statistical analyses for CM cultures were performed using the Student’s t-test on data from 3 independent experiments (n=3) with the mean of each independent experiment derived from 4 technical replicates. Analyses were done to measure differences in the CM phenotype between (1) patient and PGPC17 (healthy control) CMs (2) patient and isogenic variant-corrected CMs, (3) and patient CMs before and after drug treatment. Analyzed variables were: CM size, sarcomere organization, beat amplitude, falling time, FPA, FPD, calcium transient, phosphate concentration and protein expression. Biowire assays were analyzed using data derived from 2 cases and 2 corrected controls (3 to 13 biological replicates for each). Tukey’s multiple comparisons test was used for Biowire compaction analyses. Student’s t-test was used for nuclear area, fiber length, force, excitation threshold and maximum capture rate. Differences were considered statistically significant at p<0.05.

## Supplemental Information

Table S1: Baseline phenotype data of patient and control iPSC-CMs and cardiac Biowires

Table S2: Drug treatment data of patient iPSC-CMs and cardiac Biowires

