## Supplemental figures for "Myosin inhibitor reverses hypertrophic cardiomyopathy in pediatric iPSC-cardiomyocytes to mirror variant correction"

A

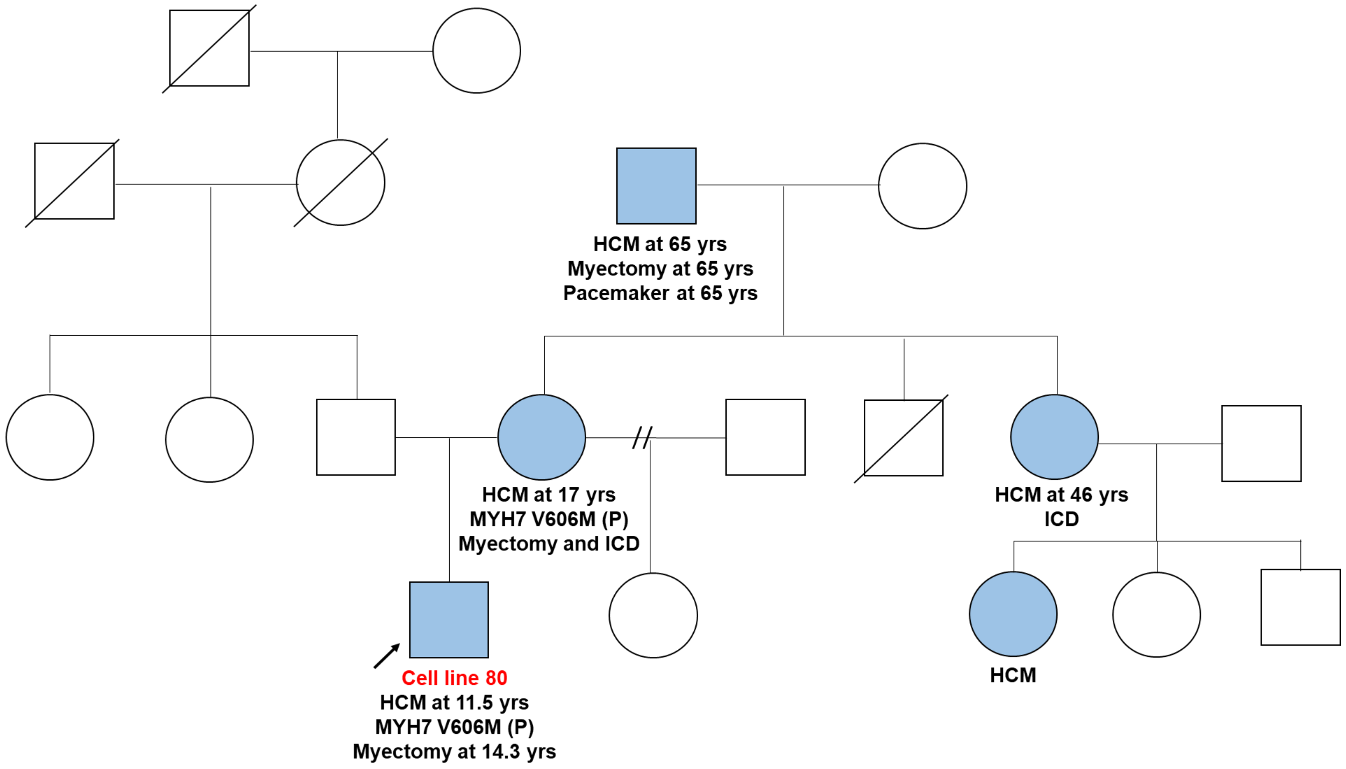

HCM at 65 yrs
Myectomy at 65 yrs
Pacemaker at 65 yrs
HCM at 17 yrs
MYH7 V606M (P)
Myectomy and ICD
HCM at 46 yrs
ICD
Cell line 80
HCM at 11.5 yrs
MYH7 V606M (P)
Myectomy at 14.3 yrs
HCM


B

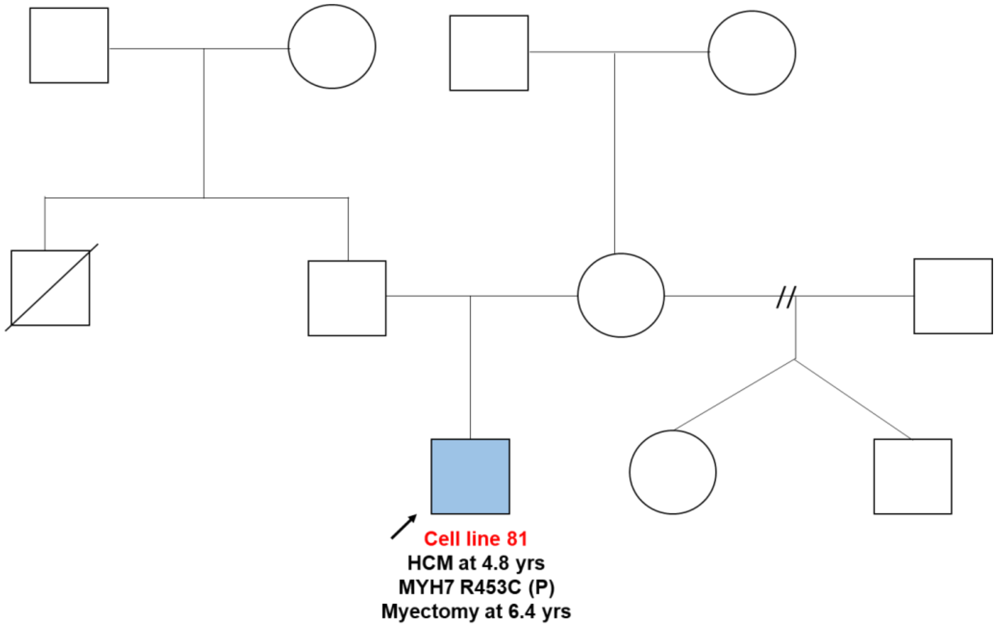

Cell line 81
HCM at 4.8 yrs
MYH7 R453C (P)
Myectomy at 6.4 yrs

C

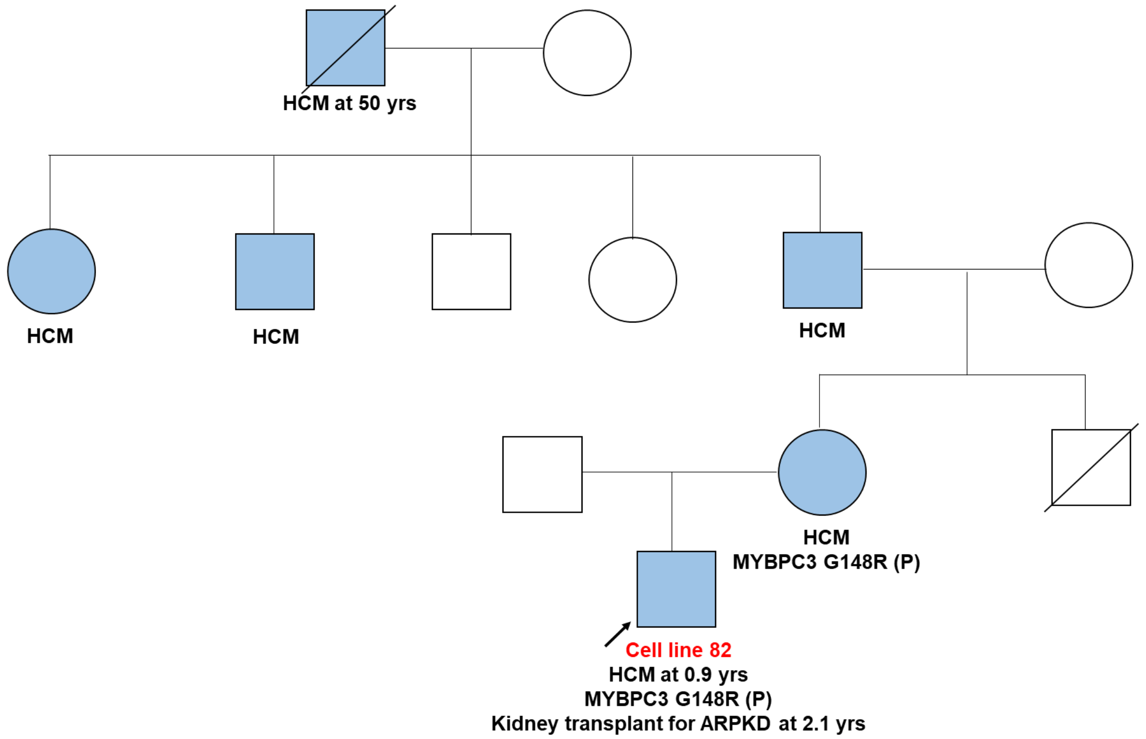


D

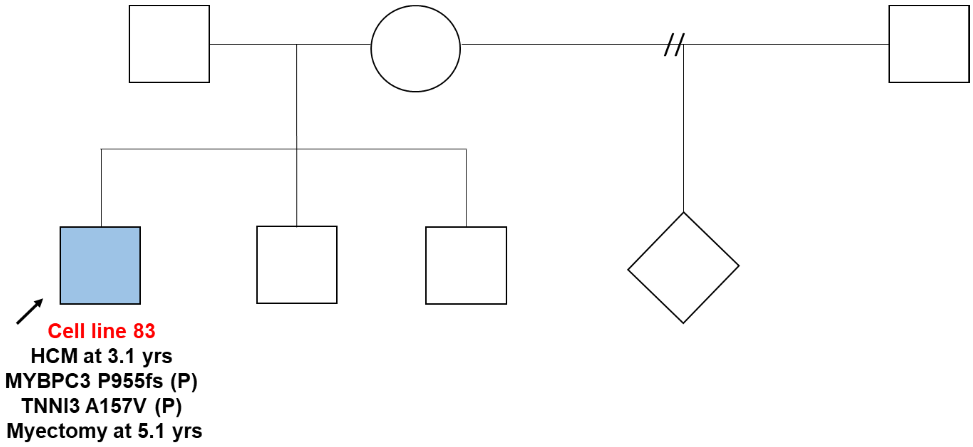


**Figure S1. Family pedigrees. A-D** Family pedigree of affected HCM probands. Circle = female, square = male; blue = affected individuals. Arrow = proband; oblique line = deceased; double lines = consanguineous. P, pathogenic variant; ICD, implantable cardioverter-defibrillator; ARPKD, autosomal recessive polycystic kidney disease.

**A**

80 (*MYH7* V606M)

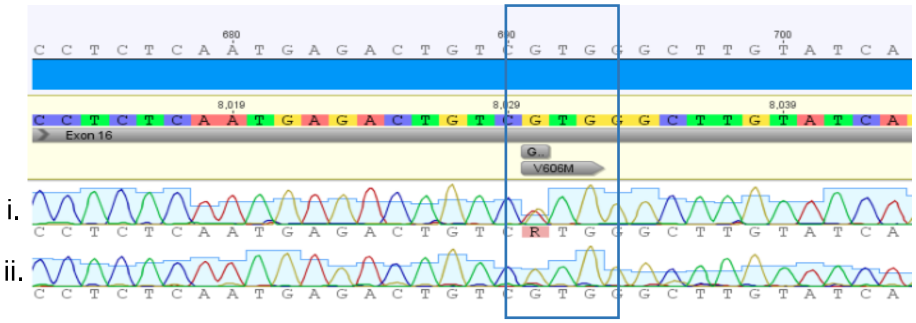


**B**

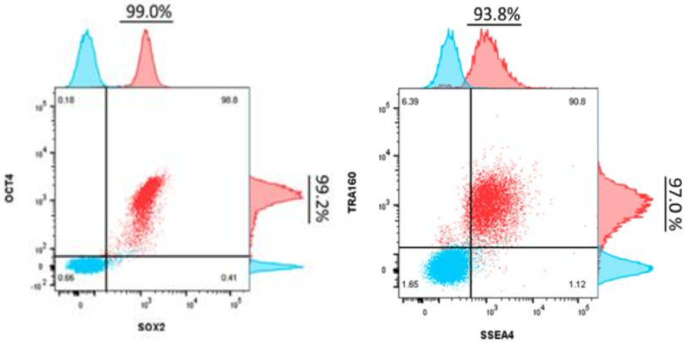


**C**

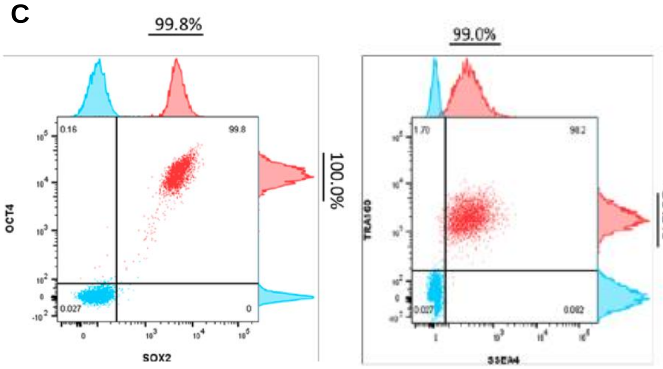


**D**

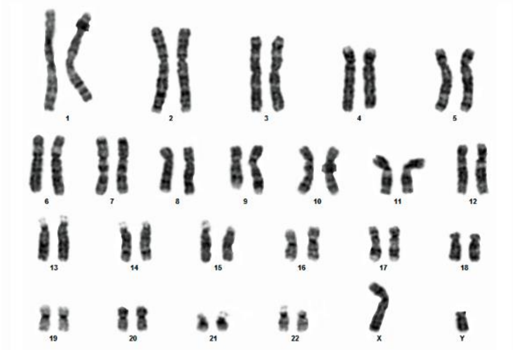

**E**

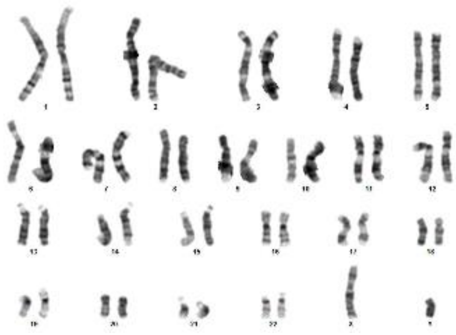

**F**

| Chromosomal co-ordinates | Potential region of cleavage | Sequencing from CCRM80V-1G |
|---|---|---|
| chr14: 23,427,653 <- 23,427,675 (MYH7 CDS) | CCTCTCAATGAGACTGTCGTGGG | CCTCTCAATGAGACTGTCGTGGG |
| chr6: 166,151,929 -> 166,151,950 | CCTTTCAATGA-ACTGTCATGGG | CCTTTCAATGAACTGTCATGGG |
| chr15: 62,911,240 <- 62,911,262 | TCTCCCACTGAGACTGTCATTGG | TCTCCCACTGAGACTGTCATTGG |
| chr17: 20,444,582 <- 20,444,606 | CCTCTCAATGCAGACTGTCCATCGG | CCTCTCAATGCAGACTGTCCATCGG |
| chr9: 104,982,256 <- 104,982,278 | CTTTTCAAAGAGACTGTCATAGG | CTTTTCAAAGAGACTGTCATAGG |

**Figure S2: Characterization of 80 patient and variant-corrected iPSCs. A.** Sequencing of *MYH7* region of interest from genomic DNA extracted from **i**) 80 parental line showing cells heterozygous for p.V606M (c.G1816A), and **ii**) 80C CRISPR edited line showing correction to homozygous wildtype. **B.** Pluripotency-associated protein expression of OCT4 and TRA160 was ≥ 90% by flow cytometry in 80 and **C.** 80C. **D.** Karyotype was normal, 46XY, for 80 and **E.** 80C. **F**. Assessment for off-target effects showed that PCR and sequencing of sites most likely to be affected by off-target cleavage of gRNA showed no off-target events for 80C.
gRNA, guide RNA.

**A**

81 (*MYH7* R453C)

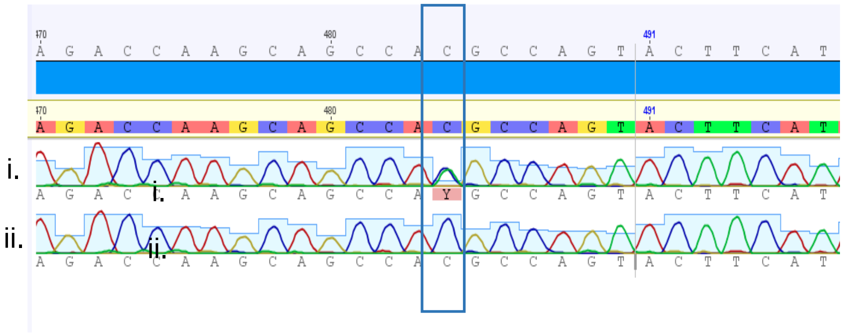


**B**

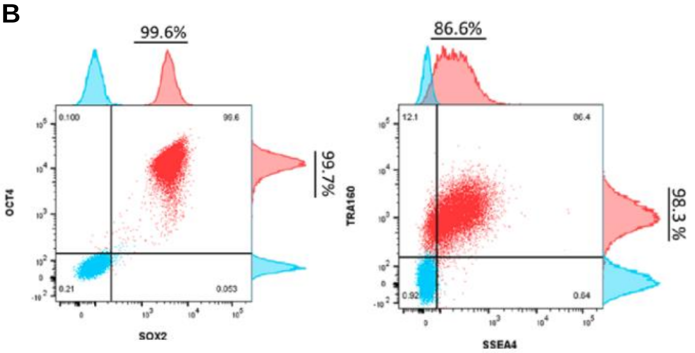


**C**

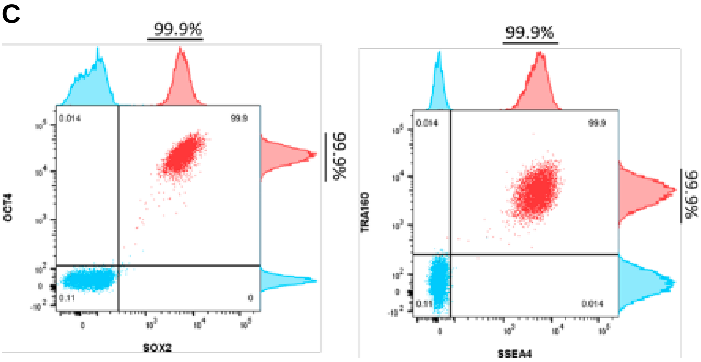


**D**

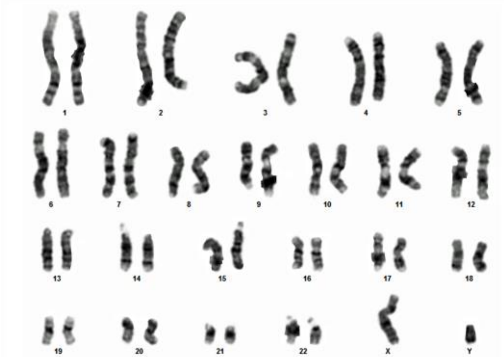

**E**

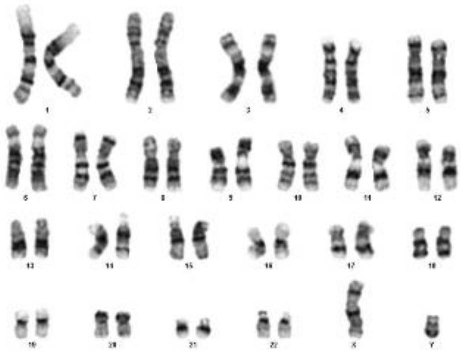

**F**

| Chromosomal co-ordinates | Potential region of cleavage | Sequence from CCRM81J-B2 |
|---|---|---|
| chr14 (NC_000014) 23,400,740 -> 23,400,762 (MYH6 CDS) | ACTCCTATGAAGTACTGGCGTGG | ACTCCTATGAAGTACTGGCGTGG |
| chr14 (NC_000014) 23,428,986 -> 23,429,008 (MYH7 CDS) | ACTCCTATGAAGTACTGGCGTGG | ACTCCTATGAAGTACTGGCGTGG |
| chr13 (NC_000013) 41,194,846 <- 41,194,868 | AACCCTATGAAATACTGGCAGGG | AACCCTATGAAATACTGGCAGGG |
| chr15 (NC_000015) 48,129,903 -> 48,129,926 | TCTCCTATCAAGTTACTGGCAGGG | TCTCCTATCAAGTTACTGGCAGGG |
| chr8 (NC_000008) 22,832,707 -> 22,832,728 | AGTCCTA-GAAGCACTGGCAAGG | AGTCCTAGAAGCACTGGCAAGG |

**Figure S3: Characterization of 81 patient and variant-corrected iPSCs. A.** Sequencing of *MYH7* region of interest from genomic DNA extracted from **i**) 81 parental line showing cells heterozygous for p.R453C (c.C1357T) and **ii**) 81C CRISPR edited line showing correction to homozygous wildtype. **B.** Pluripotency-associated protein expression of OCT4 and TRA160 was ≥ 90% by flow cytometry in 80 and **C.** 81C. **D.** Karyotype was normal, 46XY, for 81 and **E.** 81C. **F**. Assessment for off-target effects showed that PCR and sequencing of sites most likely to be affected by off-target cleavage of gRNA showed no off-target events for 81C.
gRNA, guide RNA.

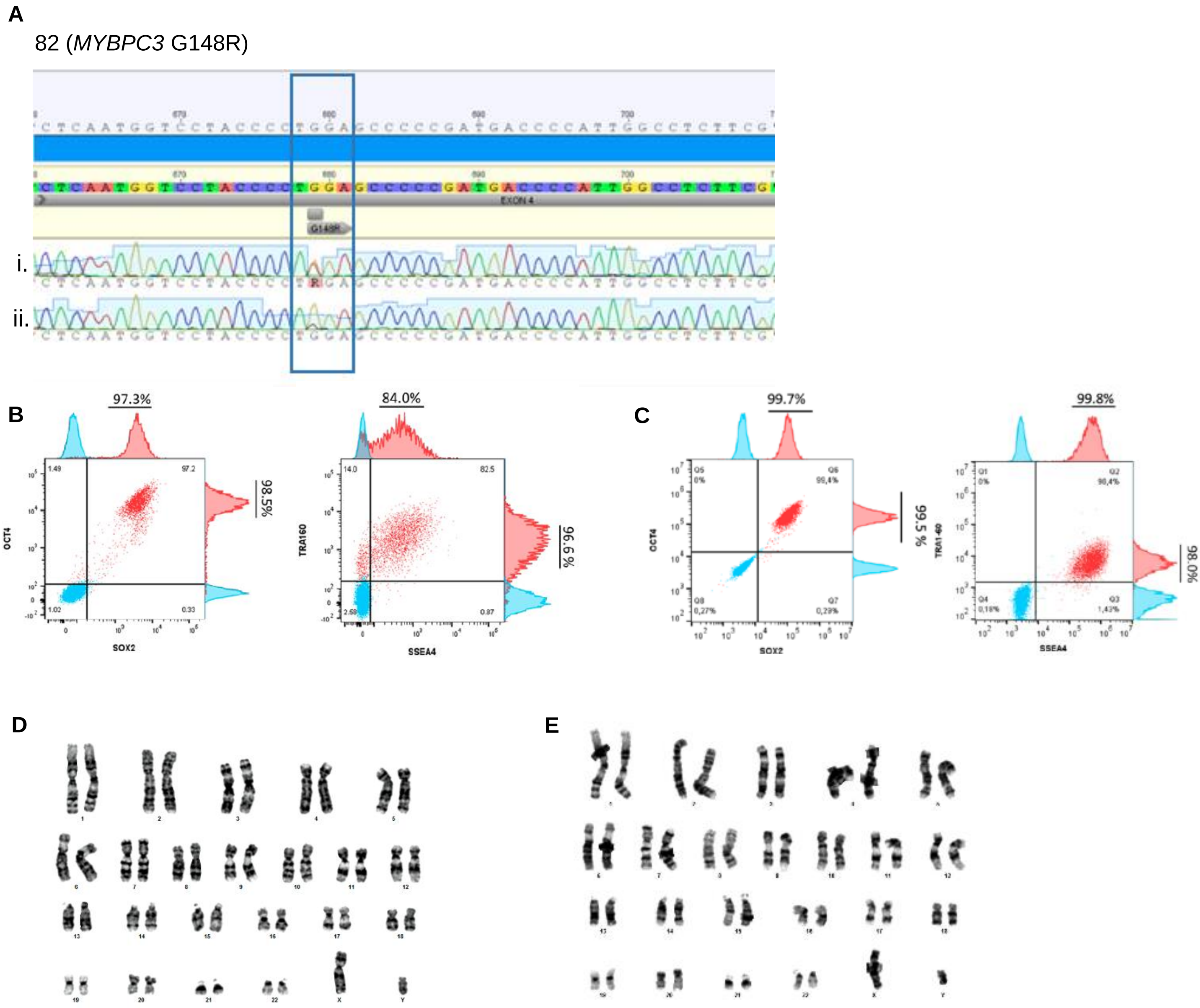


F

| Chromosomal co-ordinates | Potential region of cleavage | Sequence from CCRM82U-4B |
|---|---|---|
| chr11: 47,350,060 -> 47,350,082 (MYBPC3 CDS) | GGGGTCATCGGGGGCTCCAGGGG | GGGGTCATCGGGGGCTCCAGGGG |
| chr1: 55,533,112 <- 55,533,132 | GGGGTCA-CGGGGGCT-TAGTGG | GGGGTCACGGGGGCTTAGTGG |
| chr16: 84,451,875 <- 84,451,896 | GGGGTCACCGGGGGCT-TAGGGG | GGGGTCACCGGGGGCTTAGGGG |
| chr2: 221,572,480 -> 221,572,502 | GCGGTCTCCCGGGGCTCTAGGGG | GCGGTCTCCCGGGGCTCTAGGGG |

**Figure S4: Characterization of 82 patient and variant-corrected iPSCs. A.** Sequencing of *MYBPC3* region of interest from genomic DNA extracted from **i**) 82 parental line showing cells heterozygous for c.G442A (p.G148R), and **ii**) 82C CRISPR edited line showing correction to homozygous wildtype. **B**. Pluripotency-associated protein expression of OCT4 and TRA160 was ≥ 90% by flow cytometry in 82 and **C.** 82C. **D.** Karytoype was normal, 46XY, for 82 and **E**. 82C. **G, H.** Assessment for off-target effects showed that PCR and sequencing of sites most likely to be affected by off-target cleavage of gRNA showed no off-target events for 82C. gRNA, guide RNA.

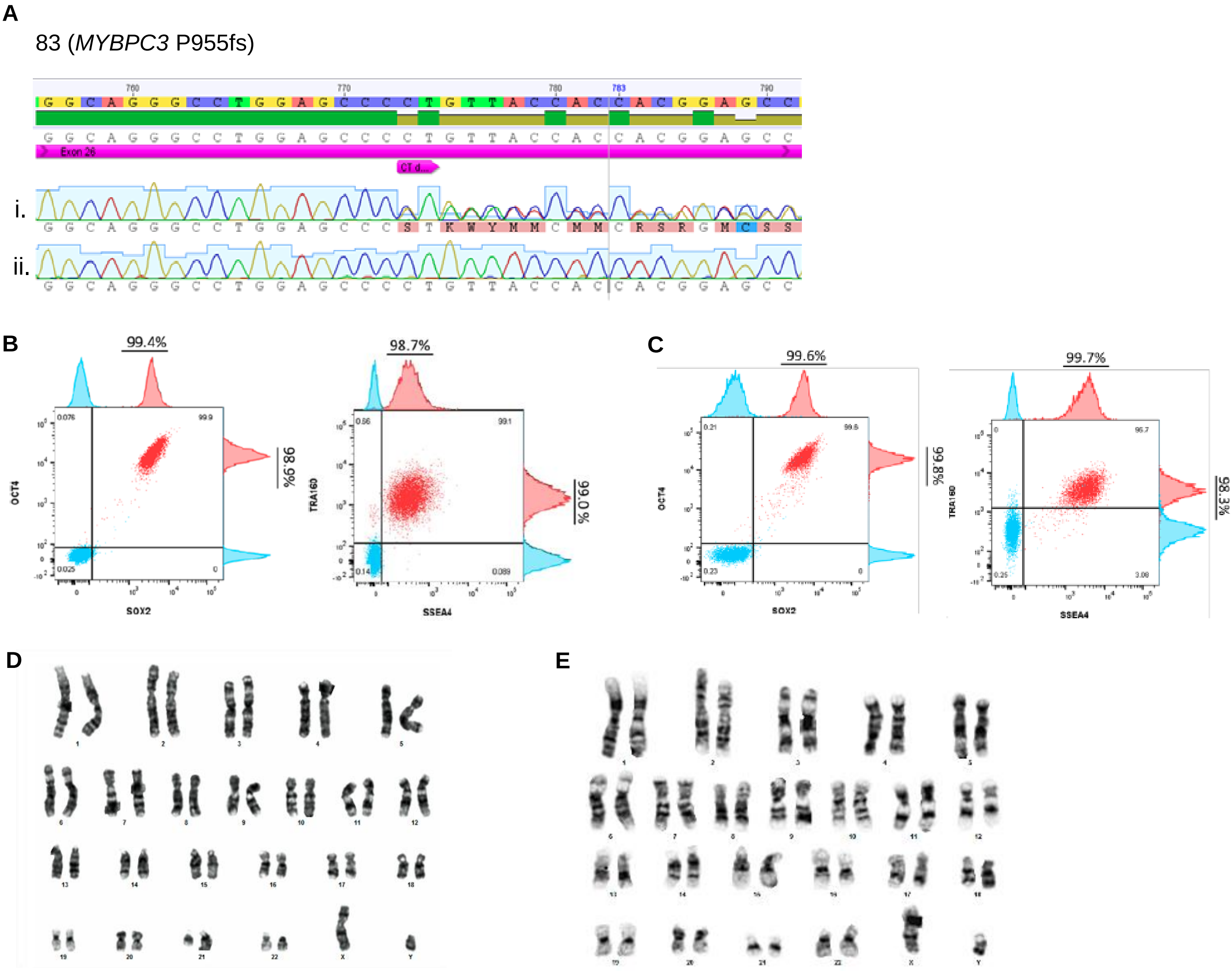


**F**

| Chromosomal co-ordinates | Potential region of cleavage | Sequence from CCRM83M-B4-E1 |
|---|---|---|
| chr16:22,723,195 - 22,723,216 | CACCGGCCCCG-GGTGGTAAGGG | CACCGGCCCCGGGTGGTAAGGG |
| chr7: 157,856,457 - 157,856,478 | CACCGGC-CTGTGGTGGGAACGG | CACCGGCCTGTGGTGGGAACGG |
| chr15: 89,784,335 - 89,784,357 | CACCGTCTTCCGTGGTGG-AAGGG | CACCGTCTTCCGTGGTGGAAGGG |
| chr18: 46,756,770 - 46,756,791 | CACCGGCGCC-TGGAGGTAAAGG | CACCGGCGCCTGGAGGTAAAGG |
| chr22: 37,918,904 - 37,918,927 | CACCAGGCTTCGTGGTGGAAATGG | CACCAGGCTTCGTGGTGGAAATGG |

**Figure S5: Characterization of 83 patient and variant-corrected iPSCs. A.** Sequencing of *MYBPC3* region of interest from genomic DNA extracted from **i**) 82 parental line showing cells heterozygous for p.P955fs (c.2864_2865del) and **ii**) 83C CRISPR edited line showing correction to homozygous wildtype. **B**. Pluripotency-associated protein expression of OCT4 and TRA160 was ≥ 90% by flow cytometry in 83 and **C.** 83C. **D.** Karytoype was normal, 46XY, for 83 and **E**. 83C. **G, H**. Assessment for off-target effects showed that PCR and sequencing of sites most likely to be affected by off-target cleavage of gRNA showed no off-target events for 83C.
gRNA, guide RNA.

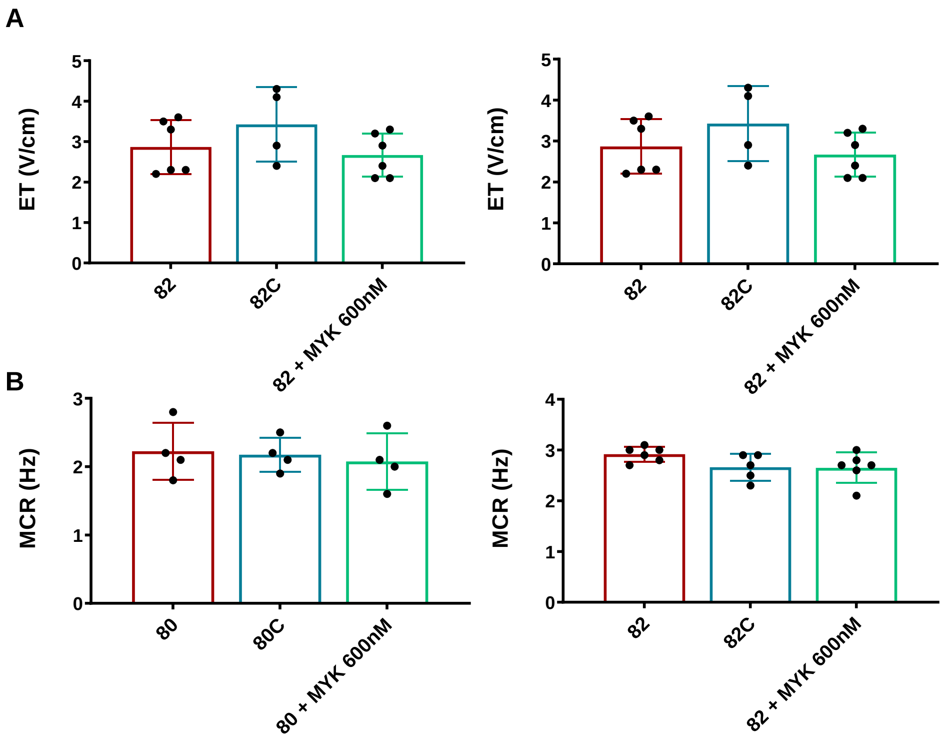


**Figure S6. Electrical excitability and calcium transients in cardiac Biowires from 80 and 82 patient iPSC-CMs. A. Excitation threshold-minimum** applied voltage required to initiate contraction was not different between patient, variant-corrected and MYK-461 treated Biowires**. B. Maximum capture rate** was not different between patient, variant-corrected and MYK-461 treated Biowires. n=4 to 6 biological replicates. Error bars represent standard deviation.
80, *MYH7* V606M; 82, *MYBPC3* G148R.
ET, excitation threshold; MCR, maximum capture rate.
